# MYC overexpression drives poor prognosis and differential sensitivity to treatments according to TP53 status in chronic lymphocytic leukemia

**DOI:** 10.64898/2026.05.14.724995

**Authors:** Alba Garrote-de-Barros, Juan Pérez-Fernández, Andrés Arroyo-Barea, Irene Bragado-García, Roberto García-Vicente, Raquel Ancos-Pintado, María Velasco-Estévez, María Linares, Joaquín Martínez-López, María Hernández-Sánchez

**Author notes:** **Corresponding author:** María Hernández-Sánchez. Department of Biochemistry and Molecular Biology. Pharmacy School, Universidad Complutense de Madrid. Plaza de Ramón y Cajal, s/n, 28040 Madrid (Spain). These authors contributed equally to this work as co-senior authors (María Linares, Joaquín Martínez-López, María Hernández-Sánchez).

## Abstract

Chronic lymphocytic leukemia (CLL) is a lymphoid neoplasm with very heterogeneous clinical and biological behavior. Among molecular variables, *TP53* alterations are well-established adverse prognostic markers; however, MYC activation, which has been linked to disease progression, has not been completely defined in terms of clinical and biological impact, particularly in relation to *TP53* status. Here, we investigated the effects of *MYC* overexpression according to *TP53* status using clinical and transcriptomic data from CLL patients and novel cellular models. CLL patients with *TP53^WT^* and *MYC* overexpression exhibited significantly shorter time to first treatment and overall survival, indicating an aggressive disease course comparable to that of patients with TP53 alterations. Consistently, *MYC* overexpression in *in vitro TP53^WT^*models was associated with increased proliferation, enrichment of AKT/mTOR signaling and upregulation of genes involved in leukemogenesis and tumor progression such as *FOXO6*. Moreover, MYC overexpression was associated with increased sensitivity to venetoclax in *TP53^WT^* cells. By contrast, the concurrence of *MYC* overexpression and *TP53* dysfunction conferred resistance to conventional CLL therapies such as BCL2 or BTK inhibitors. Of note, we identified a glycolysis inhibitor, in monotherapy or combined with BKT inhibitors, as a potential therapeutic strategy for CLL patients harboring *MYC* overexpression and *TP53* alterations.

## Introduction

CLL is a mature CD5⁺ B-cell neoplasm accumulating in blood, bone marrow, lymph nodes, and spleen (1). It represents the most common adult leukemia in Western populations with a median age around 70 years and a male predominance (1). As global populations continue to age, the burden of CLL is expected to rise significantly in the near future (2). Since CLL is a heterogeneous malignancy (3), there have been numerous efforts to identify new prognostic biomarkers to refine risk stratification and disease course (4). In parallel, the therapeutic landscape of CLL has undergone a dramatic transformation over the past two decades by the introduction of highly effective targeted therapies based on Bruton tyrosine kinase inhibitors (BTKi) and BCL-2 inhibitors (5,6). Consequently, there is a growing need to identify novel predictive biomarkers to guide treatment selection in the era of targeted therapies.

Among established molecular biomarkers in CLL, apart from IGHV mutational status (7), the presence of *TP53* alterations -including deletions on chromosome 17p (del(17p)) and/or *TP53* mutations-represent one of the strongest predictors of poor prognosis and treatment resistance (8). Whereas del(17p) appears in 4-10% of CLL patients (9,10), *TP53* can be mutated in approximately 10% of CLL at diagnosis (11) and in 80% of CLL cases with del(17p) which provokes a biallelic inactivation of *TP53* in these cases. In addition, *TP53* alterations also represent high-risk factors of Richter’s transformation (RT) which is the progression of CLL to aggressive lymphoma (12,13).

Other genetic alterations associated with poor prognosis are those involving *MYC* gene, mainly the gain of 8q (gain8q) (14–16). This chromosomal alteration has been enriched in CLL with *TP53* loss (17,18). Although genetic aberrations of *MYC* are infrequent in CLL (19), its activation plays a significant role in CLL progression and RT development (20,21). In addition, *MYC* overexpression can be due to not only to 8q-related genetic alterations but also to the presence of other driver gene mutations such as those involved in NOTCH1 signaling (22,23). However, the potential clinical impact and the biological effects of MYC activation are still not fully understood in CLL.

In this study, we aim to investigate the clinical and biological impact of *MYC* overexpression in CLL according to the status of *TP53* gene. By integrating molecular and clinical data, we seek to determine whether *MYC* overexpression may serve as a complementary biomarker to refine risk stratification in CLL and improve therapeutic decision-making in this disease. Interestingly, we define MYC expression as an independent prognostic factor, especially in CLL without *TP53* alterations that could be related with AKT/mTOR activation and high *FOXO6* expression. In addition, although the concurrence of *MYC* overexpression and *TP53* dysfunction confers drug resistance to current approved CLL treatments, we identify a glycolysis inhibitor in monotherapy or in combination with BTKi as a promising therapeutic approach for CLL with *MYC* overexpression independently of *TP53* status.

## Methods

### Clinical and biological analysis from CLL map portal data

Raw gene expression count data and corresponding clinical metadata from patients with CLL were obtained from the CLL Map Portal, a multicenter database comprising samples from multiple research institutions (11). Only patients with available RNA-seq data prior to treatment initiation and clinical and genomic data were included. In addition, seven patients were excluded following the CLL Map Portal recommendations, resulting in a cohort of 531 CLL patients. Batch effects associated with institution were corrected using the *removeBatchEffect* function from the limma package (24).

CLL patients were stratified according to *MYC* expression (*MYC^high^* or *MYC^low^*) from RNA-seq data using the maximally selected rank statistics (maxstat) method (25,26), and to the absence or presence of *TP53* genetic alterations (deletion and/or mutation) (*TP53*^altered^).

Differential expression analysis was conducted using the DESeq2 R package (v.1.46.0) (27). To account for inter-study variability, the statistical model included cohort as a covariate within the design formula (design = ∼ cohort + condition). Low-count genes were pre-filtered using filterByExpr from edgeR (v.4.0.16) (28). This filtering step retained 26725 out of 57242 genes. Significance was assessed using the Wald test, and P-values were adjusted using the Benjamini-Hochberg procedure. Differentially expressed genes (DEGs) were identified based on an absolute log2 fold change > 1.5 and an adjusted P-value < 0.05. For subsequent visualization purposes, log2 fold change shrinkage was applied using the ashr algorithm to provide more stable estimates for genes with low information or high dispersion (29).

### CLL cell lines

The human CLL cell lines PGA-1 and MEC-1 were acquired from DMSZ (The Leibniz Institute DSMZ-German Collection of Microorganisms and Cell Cultures GmbH). PGA-1 cells were grown in RPMI 1640 medium (#21875-034, Gibco) with the addition of 10% Fetal Bovine Serum (FBS) (#011-90005M, Gibco) and 1% penicillin/streptomycin (P/S) (#15140-122, Gibco). MEC-1 cells were maintained in Iscove’s MDM medium (#12440-053, Gibco), also supplemented with 10% FBS and 1% P/S. HEK 293T cells, sourced from DMSZ, were cultured in DMEM (#10566-016, Gibco) with 10% FBS and 1% P/S. All cell lines were cultured at 37°C and 5% CO2. Regular mycoplasma testing was conducted on all cell lines using DreamTaq SuperMix (#K1082, Thermo Scientific) ensuring that only mycoplasma-free cells were utilized in all experiments.

### Generation of *MYC*-overexpressing PGA-1 and MEC-1 by CRISPR/SAM

To generate CLL cellular models with *MYC* overexpression (*MYC*^OE^), PGA-1 and MEC-1 were transduced with lentiviral particles containing the plasmids lentiMPHv2 (#89308, Addgene) and lentiSAMv2 (#75112, Addgene) containing the sgRNAs to overexpress endogenous *MYC* gene. These sgRNAs were designed using the CRISPick tool (30,31), to target the *MYC* promoter. Some of the selected sgRNAs were also previously reported (32). To mitigate possible biases due to off-target effects of the sgRNAs, *MYC*^OE^ CLL models were generated using different sgRNAs. Five sgRNAs were selected for downstream experiments (19, 173, 178, 240, 369) and the three best ones to induce *MYC*^OE^ were selected for functional analysis (Table S1). The plasmid lentiSAMv2 without any sgRNAs was used as a control. The selected sgRNAs were cloned into the lentiSAMv2 vector as previously described (33,34). The correct insertion of the sgRNAs was confirmed by Sanger sequencing.

Lentiviral particle production was performed as previously described (35). Lentivirus were used to infect 5 × 10^5^ PGA-1 or MEC-1 cells in medium supplemented with 8 µg/mL polybrene (Santa Cruz, sc-134220). Subsequently, cells were selected with hygromycin (300 µg/mL for both cell lines) and blasticidin (8 µg/mL for MEC-1 and 6 µg/mL for PGA-1) for 7 days.

### Generation of CLL cell lines with *TP53* knockdown and *MYC* overexpression

To induce *TP53* knockdown (*TP53^KD^*) and *MYC* overexpression (*MYC^cOE^*) in the same cell line, PGA-1 cells were transduced with lentivirus expressing vectors with *TP53*-targeting shRNA (PLKO-p53-shRNA941, Addgene #25637) and/or exogenous *MYC* cDNA (PLV-Bsd-hMYC, Vector Builder). Control cells were transduced with the corresponding empty vector (PLKO-1, Addgene #8453; PLV-Bsd-ORF, Vector Builder). Following transduction, cells were selected with puromycin (#A1113803, Gibco) (0.25 µg/mL) for shRNA constructs or blasticidin (#A1113903, Gibco) (6 µg/mL) for cDNA constructs for 7 days.

### RNA extraction of cellular models

Total RNA and were extracted from the previously generated CRISPR/SAM and shRNA/cDNA cell line models, including their corresponding biological replicates. RNA and DNA were isolated using the AllPrep DNA/RNA Kit (#80204, Qiagen) according to the manufacturer’s instructions.

### RNA-seq in CRISPR/SAM cell lines

RNA sequencing (RNA-seq) was performed on the CLL-derived cell lines PGA-1 and MEC-1 under *MYC* wild-type (*MYC*^WT^) and *MYC* overexpression (*MYC*^OE^) conditions, with three biological replicates per condition (3 different sgRNAs). Library preparation was carried out using the QuantSeq 3‘ mRNA-Seq V2 Library Prep Kit (FWD) for Illumina (catalog no. 191, Lexogen) and sequenced on a NextSeq 550 platform (RRID:SCR_016381; Illumina). This analysis was carried out externally at the Genomic Unit of the Spanish National Cancer Research Center (CNIO).

Raw sequencing reads in FASTQ format were subjected to quality control using FastQC and aggregated with MultiQC (36). Adapter trimming and poly(A) removal, as well as UMI extraction, were performed using BBDuk and UMI-tools (37,38). Read origin was validated using FastQ Screen (37). High-quality reads were aligned to the human reference genome (GRCh38) using the STAR aligner (39). UMI-based deduplication was performed, and gene-level count matrices were generated using FeatureCounts (40).

Principal component analysis (PCA) was performed on VST-transformed data to assess global transcriptional differences in both patients and cell lines.

Differential gene expression (DGE) analysis was performed using DESeq2 (27), comparing *MYC*^OE^ versus *MYC*^WT^ conditions independently in each cell line (PGA-1 and MEC-1). p-value< 0.05 and an absolute log2FC≥ 0.58 were considered significantly differentially expressed following the same strategy than we specified in patient section.

Gene set enrichment analysis (GSEA) was performed on ranked gene lists derived from differential expression analyses for cell line datasets. GSEA was performed comparing *MYC^OE^*versus *MYC^WT^* conditions in each cellular model.

Analyses were carried out using GSEA software (version 4.3.2) (41) with 1,000 permutations and the Hallmark gene set collection as reference. Pathways with a significant normalized enrichment score (NES), false discovery rate (FDR) < 0.25, and adjusted p-value < 0.05 were considered biologically relevant. Detailed bioinformatic analysis was described in Supplementary Methods.

### RT-qPCR

RT-qPCR was performed to validate *MYC* expression in the cellular models as well as to analyze the most interesting deregulated genes from transcriptomic analysis. First, cDNA was synthesized from 500 ng of total RNA using the High-Capacity cDNA Reverse Transcription Kit (#4374967, Applied Biosystems), following the manufacturer’s instruction Quantitative real-time PCR was performed using FastStart Universal SYBR Green Master (ROX) (#4913850001, Roche). Relative gene expression levels were calculated using the comparative ΔΔCt method, with GAPDH used as the endogenous reference gene. Supplementary Table 2 shows the oligo sequences of the analyzed genes (Table S2).

### Western blot

Cells were resuspended in 1× RIPA lysis buffer (#20-188, Millipore,) supplemented with protease and phosphatase inhibitors (#4693124001, #4906845001, Roche). Protein concentration was determined using the Pierce™ BCA Protein Assay Kit (#A55864, Thermo Scientific^TM^). Protein samples were mixed loading buffer (#1610747, Bio-Rad) supplemented with β-mercaptoethanol and denatured at 95 °C prior to electrophoresis.

Denatured protein samples, together with a molecular weight marker (##1610374, Biorad), were separated by SDS–PAGE using the Mini-PROTEAN® Tetra Cell 4-Gel System (Bio-Rad) and transferred onto PVDF membranes (#1620184, Bio-Rad), which were previously activated with methanol.

Following transfer, membranes were blocked with 3% BSA and subsequently incubated with the appropriate primary antibodies overnight at 4 °C (Table S3). After overnight incubation with primary antibodies, membranes were washed several times with TBS-T and subsequently incubated with the appropriate anti-rabbit secondary antibody (Table S3). Protein signals were detected using either the Cytiva ECL Start Kit (# RPN2209, Cytiva) for total protein and the SuperSignal™ West Femto Maximum Sensitivity Substrate (#34095, Thermo Scientific^TM^) for phosphorylated proteins, according to the manufacturer’s instructions. Chemiluminescent signals were captured using the ChemiDoc™ MP Imaging System (Bio-Rad).

### Inmunofluorescence

500.000 cells per condition were fixed with 8% PFA in PBS for 10 minutes, permeabilized and blocked with blocking buffer (0.5% BSA, 0.1% Triton X-100 in 1X PBS) overnight at 4°C. After this, the samples were cleaned with PBS and incubated with 1:100 dilution of MYC primary antibody (#ab190560, Abcam) in blocking buffer, overnight at 4°C. After washing with PBS, the secondary antibody, Alexa FluorTM 647 donkey anti-rabbit IgG (1:500, #A-32795, Invitrogen), was added for 90 minutes in BSA. After washing with PBS, cell nuclei were stained with 10 µg/mL DAPI (#D1306, Invitrogen) for 15mins. Images were acquired using a Leica SP5 confocal microscope.

### Cell proliferation assay

Cell proliferation was assessed for CRISPR/SAM and cDNA/shRNA cellular models using a 96-well plate format. Cells were seeded in triplicate at a density of 20,000 cells/ml. After 3 and 6 days, WST-8 reagent from the Cell Counting Kit-8 Reagent (#96992, Sigma-Aldrich) was added to each well. Plates were incubated for 2 h at 37 °C, and absorbance was measured at 450 nm and 620 nm for wavelength correction, using a Gen5 2.0 software (BioTek Instruments) plate reader. Proliferation rates were calculated as the mean absorbance for each experimental condition across the triplicates.

### Cell Cycle analysis

Cell cycle distribution was analyzed by propidium iodide (PI) staining using the FxCycle™ PI/RNase Staining Solution (#F10797, Invitrogen), according to the manufacturer’s instructions. Briefly, 1 × 10^6^ cells per condition were cultured and, at 48 hours, cells were washed twice with phosphate-buffered saline (PBS), and fixed by the dropwise addition of 70% cold ethanol. Following fixation, cells were incubated with FxCycle™ PI/RNase staining solution for 15–30 minutes at room temperature in the dark. Samples were acquired using a BD FACSCanto II flow cytometer (BD Biosciences) and data were analyzed using FlowJo software. This experiment was performed in three biological replicates per condition.

### Apoptosis analysis

Apoptosis was assessed using Annexin V (#640941, BioLegend) and PI staining (#P4864, Merck) followed by flow cytometry. In brief, 1 × 10^6^ cells were seeded for 48 hours and then labelled with Annexin V and PI. After staining, cells were immediately analyzed using BD FACSCanto II flow cytometer (BD Biosciences) and data were analyzed using FlowJo software. Data were analyzed to distinguish viable, early apoptotic, late apoptotic, and necrotic cell populations.

### Drug treatments and viability assays

Cell viability assays were performed in CRISPR/SAM and cDNA/shRNA cellular models after drug exposure. Drugs used were BTK inhibitor (BTKi) ibrutinib (#0111-I-3311, Gentaur), BCL-2 inhibitor (BCL-2i) venetoclax (#0111-V-3579, Gentaur) and JQ1 (#HY-13030, MedChemExpress). A library of 203 drugs was obtained from MedChemExpress (Table S4). High-throughput drug library was tested at a single drug concentration (20µM) in PGA-1 *TP53*^KD^/*MYC*^cOE^. The most promising drugs from the library and CLL-approved treatments such as ibrutinib or venetoclax were tested with increasing concentrations in cellular models with different alterations of *MYC* and *TP53*.

Cells were seeded at a density of 1.5 × 10^5^ cells/ml and treated for 72 hours, after which cell viability was assessed using WST-8 (Cell Counting Kit-8 Reagent, #96992, Sigma-Aldrich). The percentage of cell survival from three independent biological replicates per condition was calculated and normalized to DMSO-treated controls.

### Statistical analysis

Statistical analyses of experimental data were performed using GraphPad Prism software (V 8.0.1) (GraphPad Software, Inc.) and R (Version 4.4.3).

Data are presented as mean ± standard error of the mean (SEM) from at least two or three independent biological replicates, as indicated in the corresponding figure legends. Normality of the data distribution was assessed prior to statistical testing. For comparisons between two groups, an unpaired two-tailed Student’s t-test was applied for normally distributed data, whereas the non-parametric Mann–Whitney U test was used when normality was not assumed. For comparisons involving more than two groups, one-way or two-way analysis of variance (ANOVA) was performed.

Correlation between gene expression levels was assessed using Spearman’s rank correlation coefficient. Only samples with detectable expressions were included in the analysis.

Overall survival (OS) and time to first treatment (TFT) were analyzed by Kaplan–Meier curves using the *survfit* function (V.3.8.3) (42) and visualized with *ggsurvplot* (V.0.5.0) (43). Statistical significance was assessed using the log-rank test, with multiple testing correction performed using the Benjamini–Hochberg (BH) method when applicable.

Univariate and multivariable logistic regression analyses were performed to evaluate associations between biological and clinical variables with *MYC* expression levels. Cox proportional hazards regression models were applied to assess the impact of these variables on overall survival (OS) and time to first treatment (TFT). All variables were initially examined in univariate models. Variables with a p-value ≤ 0.2 in the univariate analysis, and genomic alterations present in at least 8% of patients, were considered for inclusion in multivariable models. Additionally, clinically relevant variables, including age, IGHV mutational status, and sex, were included in multivariable analyses irrespective of their univariate significance. Results are reported as odds ratios (ORs) for logistic regression and hazard ratios (HRs) for Cox models, both with 95% confidence intervals (CIs). The proportional hazards assumption was evaluated using Schoenfeld residuals.

Statistical significance was defined as p ≤ 0.05. Significance levels in figures are represented as follows: *p ≤ 0.05, **p ≤ 0.01, ***p ≤ 0.001, ****p ≤ 0.0001.

## Results

### High levels of *MYC* expression represented an independent biomarker of poor prognosis in *TP53*^WT^ CLL patients

*MYC^high^* expression was associated independently with the presence of trisomy 12 or *TP53* alterations (*TP53^altered^*) (Table 1). Interestingly, we observed that high levels of *MYC* expression (*MYC^high^*) were significantly associated with shorter TFT and OS in CLL (Figure S1A). Although MYC expression did not represent an independent biomarker in terms of TFT (Table S5), it remained as an independent predictor of poor OS in the whole cohort (Table 2). In addition, when we stratified CLL patients according to *TP53* status, we showed that *MYC^high^*was significantly associated with inferior TFT and OS in *TP53^WT^*patients with a similar progression to *TP53^altered^* group (Figures 1A–B). By contrast, *MYC* expression did not have clinical impact in CLL with *TP53^altered^*. Of note, *MYC* expression represented an independent prognostic factor for both OS and TFT in *TP53^WT^* CLL patients (Tables 3-4).

**Figure 1:**
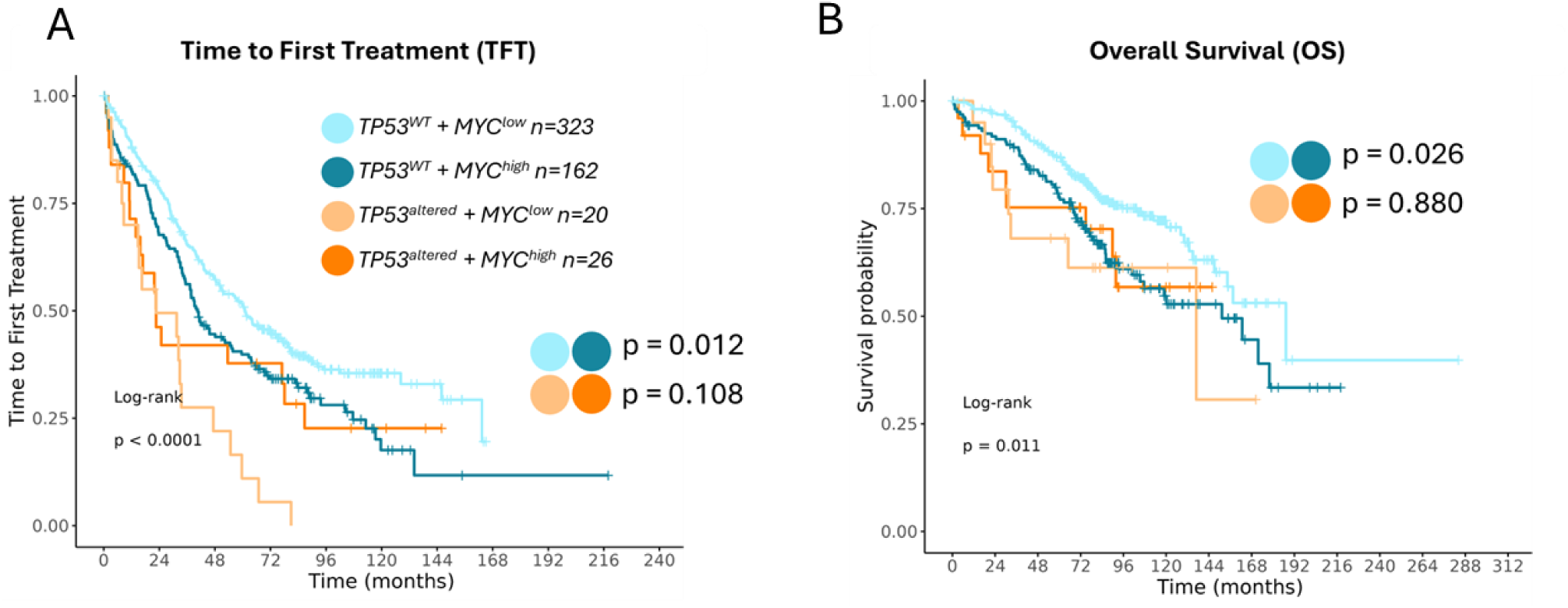
Clinical impact of *MYC* expression according to *TP53* status in CLL patients. Kaplan–Meier plots showing the association between *MYC* expression and *TP53* alterations with the time to first treatment (A) and overall survival (B) in 531 untreated CLL patients from the CLL-map portal. CLL patients were grouped according to *MYC* expression levels and TP53 status: *TP53^WT^ + MYC^low^*(light blue line), *TP53^WT^ + MYC^high^* (dark blue line), *TP53^altered^ + MYC^low^* (light orange line) and *TP53^WT^ + MYC^high^* (dark orange line).

**Table 1:**
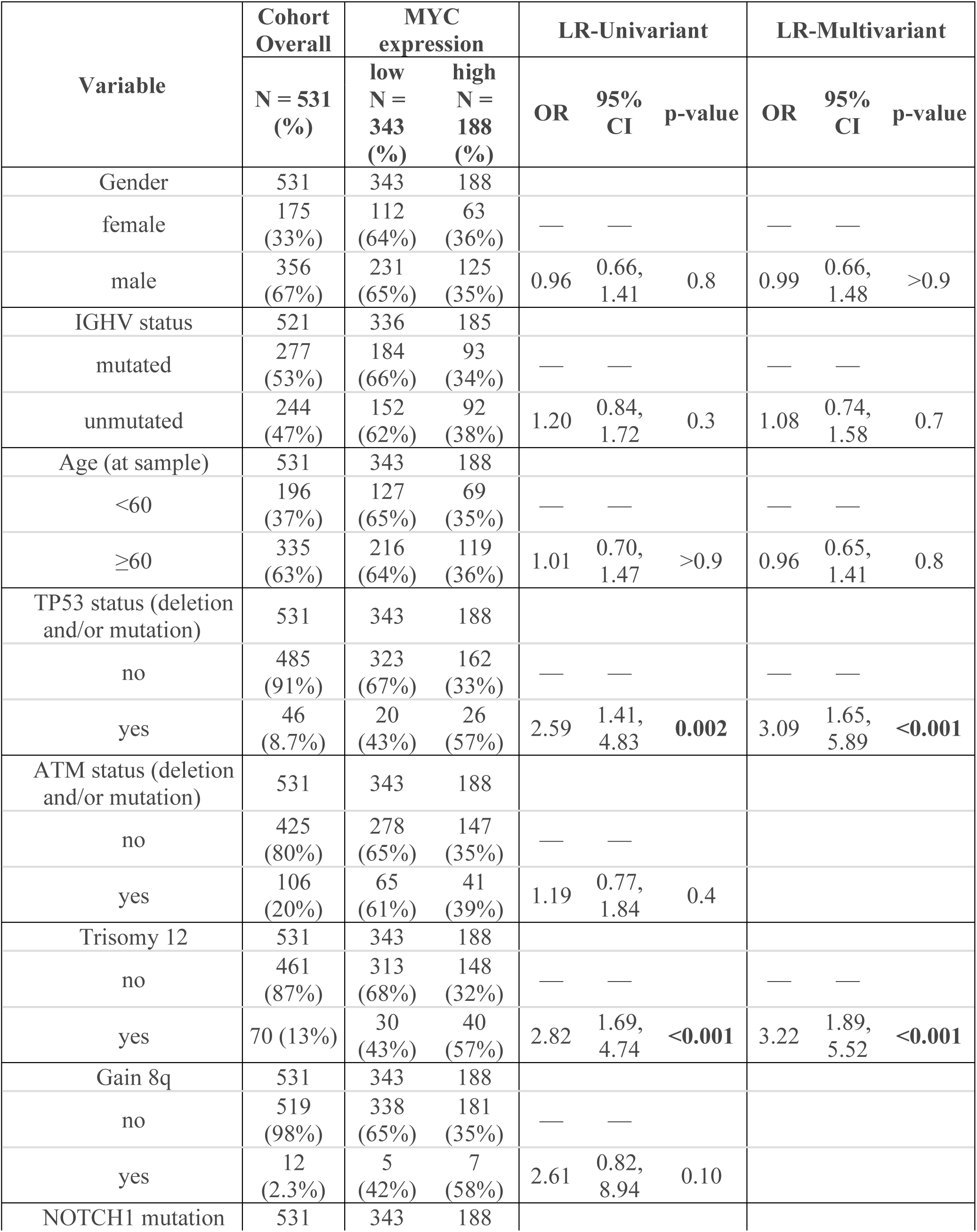
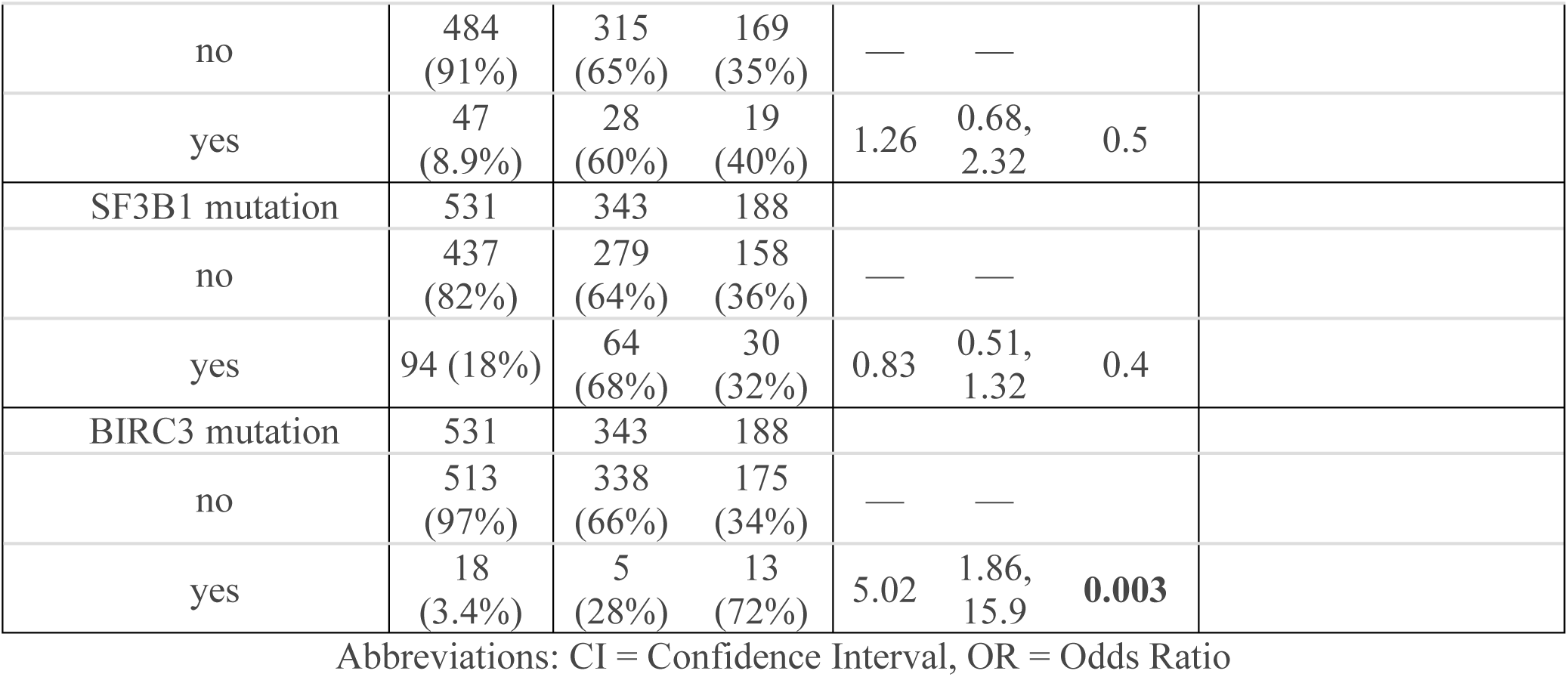
Logistic regression of 531 untreated CLL patients from CLL-map portal.

| Variable | Cohort Overall | MYC expression |  | LR-Univariant |  |  | LR-Multivariate |  |  |
| --- | --- | --- | --- | --- | --- | --- | --- | --- | --- |
|  | N = 531 (%) | low N = 343 (%) | high N = 188 (%) | OR | 95% CI | p-value | OR | 95% CI | p-value |
| Gender | 531 | 343 | 188 |  |  |  |  |  |  |
| female | 175 (33%) | 112 (64%) | 63 (36%) | — | — |  | — | — |  |
| male | 356 (67%) | 231 (65%) | 125 (35%) | 0.96 | 0.66, 1.41 | 0.8 | 0.99 | 0.66, 1.48 | >0.9 |
| IGHV status | 521 | 336 | 185 |  |  |  |  |  |  |
| mutated | 277 (53%) | 184 (66%) | 93 (34%) | — | — |  | — | — |  |
| unmutated | 244 (47%) | 152 (62%) | 92 (38%) | 1.20 | 0.84, 1.72 | 0.3 | 1.08 | 0.74, 1.58 | 0.7 |
| Age (at sample) | 531 | 343 | 188 |  |  |  |  |  |  |
| <60 | 196 (37%) | 127 (65%) | 69 (35%) | — | — |  | — | — |  |
| ≥60 | 335 (63%) | 216 (64%) | 119 (36%) | 1.01 | 0.70, 1.47 | >0.9 | 0.96 | 0.65, 1.41 | 0.8 |
| TP53 status (deletion and/or mutation) | 531 | 343 | 188 |  |  |  |  |  |  |
| no | 485 (91%) | 323 (67%) | 162 (33%) | — | — |  | — | — |  |
| yes | 46 (8.7%) | 20 (43%) | 26 (57%) | 2.59 | 1.41, 4.83 | **0.002** | 3.09 | 1.65, 5.89 | **<0.001** |
| ATM status (deletion and/or mutation) | 531 | 343 | 188 |  |  |  |  |  |  |
| no | 425 (80%) | 278 (65%) | 147 (35%) | — | — |  |  |  |  |
| yes | 106 (20%) | 65 (61%) | 41 (39%) | 1.19 | 0.77, 1.84 | 0.4 |  |  |  |
| Trisomy 12 | 531 | 343 | 188 |  |  |  |  |  |  |
| no | 461 (87%) | 313 (68%) | 148 (32%) | — | — |  | — | — |  |
| yes | 70 (13%) | 30 (43%) | 40 (57%) | 2.82 | 1.69, 4.74 | **<0.001** | 3.22 | 1.89, 5.52 | **<0.001** |
| Gain 8q | 531 | 343 | 188 |  |  |  |  |  |  |
| no | 519 (98%) | 338 (65%) | 181 (35%) | — | — |  |  |  |  |
| yes | 12 (2.3%) | 5 (42%) | 7 (58%) | 2.61 | 0.82, 8.94 | 0.10 |  |  |  |
| NOTCH1 mutation | 531 | 343 | 188 |  |  |  |  |  |  |

|  |  |  |  |  |  |  |
| --- | --- | --- | --- | --- | --- | --- |
| no | 484<br>(91%) | 315<br>(65%) | 169<br>(35%) | — | — |  |
| yes | 47<br>(8.9%) | 28<br>(60%) | 19<br>(40%) | 1.26 | 0.68,<br>2.32 | 0.5 |
| SF3B1 mutation | 531 | 343 | 188 |  |  |  |
| no | 437<br>(82%) | 279<br>(64%) | 158<br>(36%) | — | — |  |
| yes | 94 (18%) | 64<br>(68%) | 30<br>(32%) | 0.83 | 0.51,<br>1.32 | 0.4 |
| BIRC3 mutation | 531 | 343 | 188 |  |  |  |
| no | 513<br>(97%) | 338<br>(66%) | 175<br>(34%) | — | — |  |
| yes | 18<br>(3.4%) | 5<br>(28%) | 13<br>(72%) | 5.02 | 1.86,<br>15.9 | **0.003** |

Abbreviations: CI = Confidence Interval, OR = Odds Ratio

**Table 2:** COX model for overall survival of 531 untreated CLL patients from CLL-map portal.

| Variable | Cohort<br>Overall | COX-OS-Univariate |  |  | COX-OS-Multivariate |  |  |
| --- | --- | --- | --- | --- | --- | --- | --- |
|  | N = 531<br>(%) | HR | 95%<br>CI | p-value | HR | 95%<br>CI | p-value |
| Gender | 531 |  |  |  |  |  |  |
| female | 175 (33%) | — | — |  | — | — |  |
| male | 356 (67%) | 1.76 | 1.22,<br>2.55 | **0.003** | 1.41 | 0.96,<br>2.07 | 0.08 |
| IGHV status | 521 |  |  |  |  |  |  |
| mutated | 277 (53%) | — | — |  | — | — |  |
| unmutated | 244 (47%) | 3.18 | 2.26,<br>4.47 | **<0.001** | 2.77 | 1.89,<br>4.04 | **<0.001** |
| Age (at sample) | 531 |  |  |  |  |  |  |
| <60 | 196 (37%) | — | — |  | — | — |  |
| ≥60 | 335 (63%) | 1.55 | 1.11,<br>2.17 | **0.01** | 1.68 | 1.19,<br>2.38 | **0.003** |
| TP53 status (deletion<br>and/or mutation) | 531 |  |  |  |  |  |  |
| no | 485 (91%) | — | — |  | — | — |  |
| yes | 46 (8.7%) | 1.55 | 0.94,<br>2.57 | 0.088 | 1.18 | 0.70,<br>1.98 | 0.5 |
| ATM status (deletion<br>and/or mutation) | 531 |  |  |  |  |  |  |
| no | 425 (80%) | — | — |  | — | — |  |
| yes | 106 (20%) | 1.87 | 1.32,<br>2.64 | **<0.001** | 1.22 | 0.84,<br>1.77 | 0.3 |
| Trisomy 12 | 531 |  |  |  |  |  |  |
| no | 461 (87%) | — | — |  | — | — |  |
| yes | 70 (13%) | 1.32 | 0.86,<br>2.03 | 0.2 | 1.23 | 0.79,<br>1.93 | 0.4 |
| Gain 8q | 531 |  |  |  |  |  |  |
| no | 519 (98%) | — | — |  |  |  |  |
| yes | 12 (2.3%) | 2.56 | 1.25,<br>5.21 | **0.01** |  |  |  |
| NOTCH1 mutation | 531 |  |  |  |  |  |  |
| no | 484 (91%) | — | — |  | — | — |  |
| yes | 47 (8.9%) | 1.44 | 0.88,<br>2.35 | 0.14 | 0.91 | 0.55,<br>1.52 | 0.7 |
| SF3B1 mutation | 531 |  |  |  |  |  |  |
| no | 437 (82%) | — | — |  | — | — |  |
| yes | 94 (18%) | 1.88 | 1.32,<br>2.67 | **<0.001** | 1.43 | 0.98,<br>2.09 | 0.06 |
| BIRC3 mutation | 531 |  |  |  |  |  |  |
| no | 513 (97%) | — | — |  |  |  |  |
| yes | 18 (3.4%) | 2.22 | 1.17,<br>4.22 | **0.02** |  |  |  |
| MYC expression level | 531 |  |  |  |  |  |  |
| low | 343 (65%) | — | — |  | — | — |  |
| high | 188 (35%) | 1.54 | 1.13,<br>2.11 | **0.007** | 1.47 | 1.06,<br>2.03 | **0.02** |

Abbreviations: CI = Confidence Interval, HR = Hazard Ratio

**Table 3:**
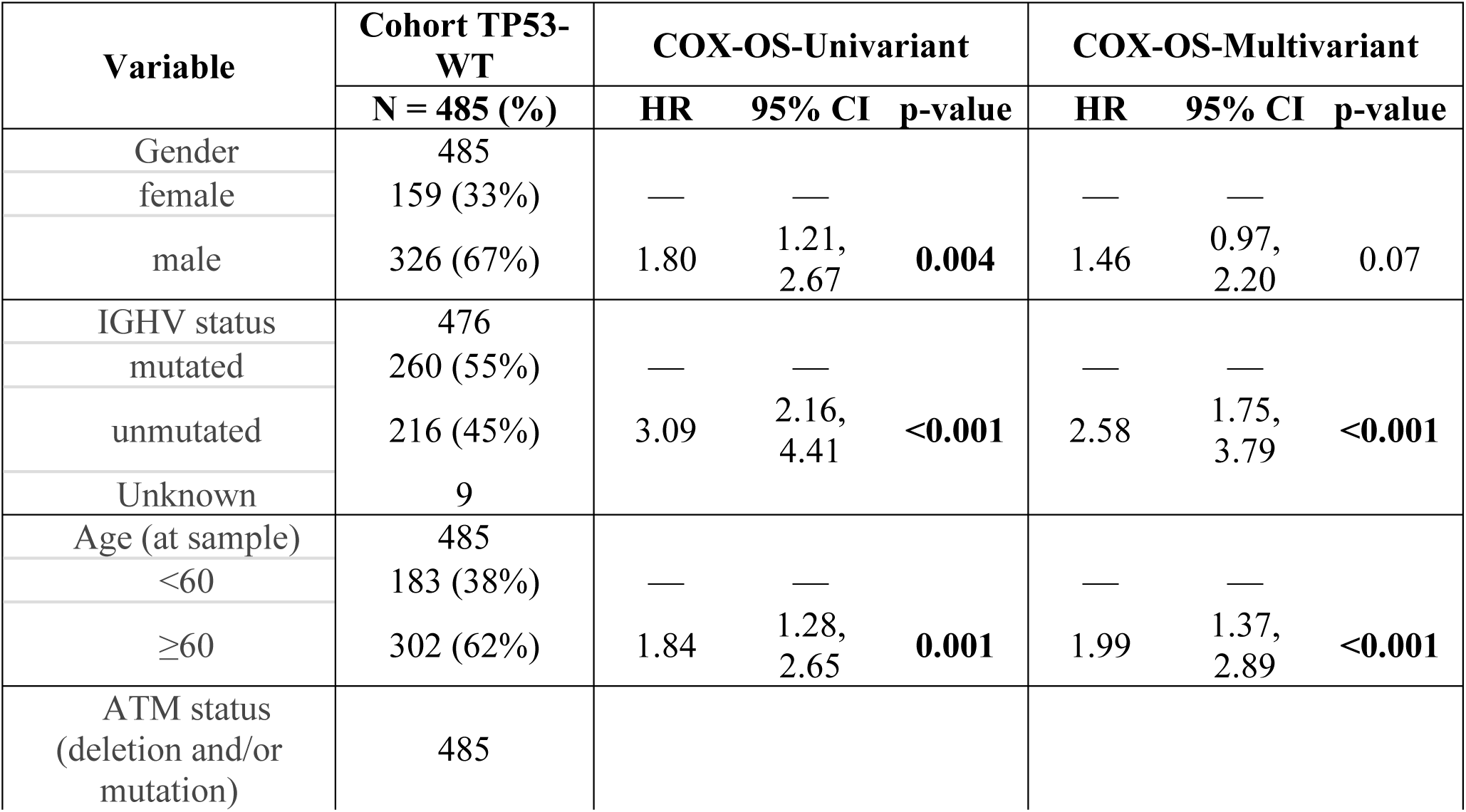
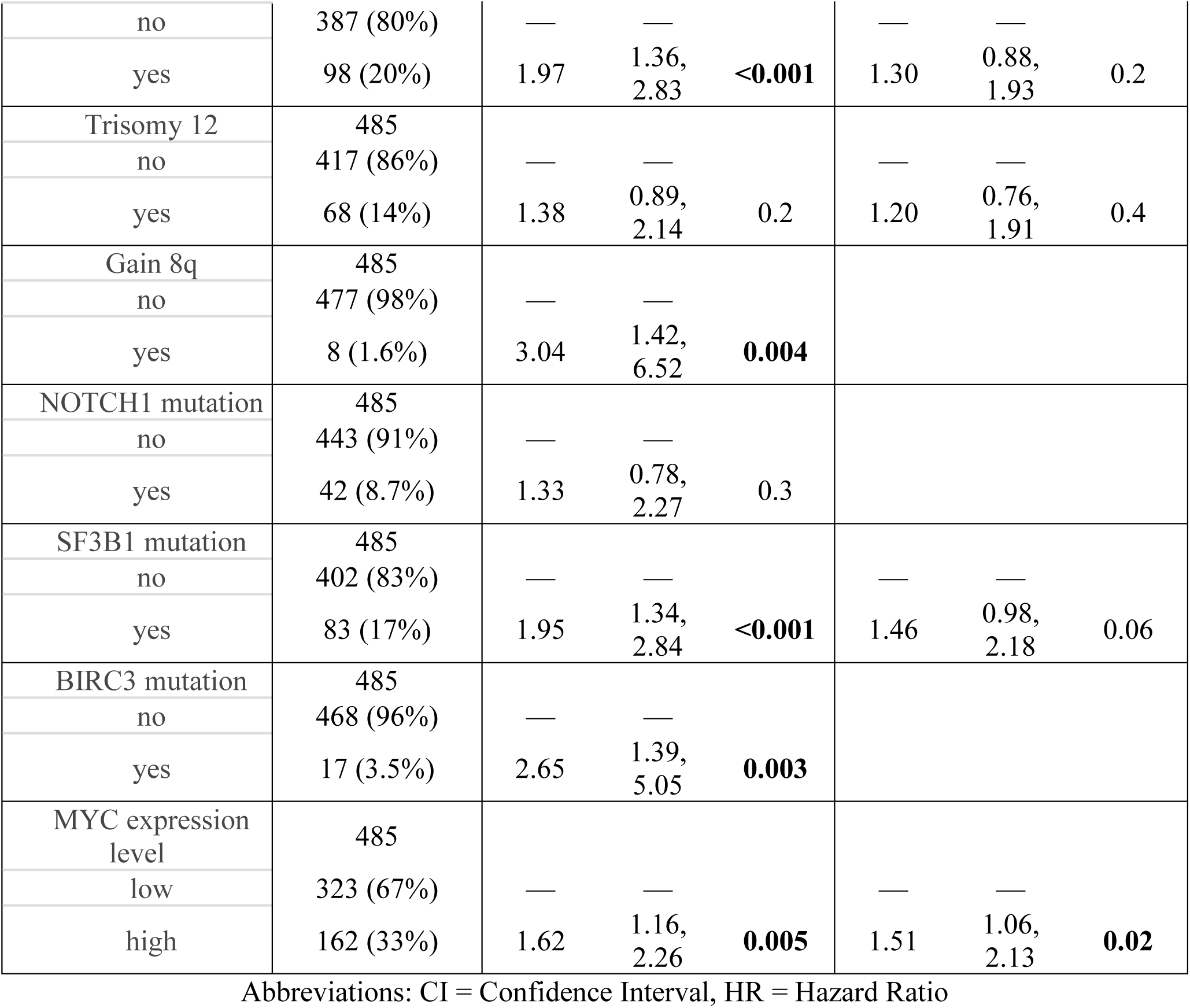
COX model for overall survival of 485 untreated *TP53^WT^* CLL patients from CLL-map portal.

**Table 4:** COX model for time to first treatment of 485 untreated *TP53^WT^* CLL patients from CLL-map portal.

| Variable | Cohort TP53-WT | COX-TFT-Univariant |  |  | COX-TFT-Multivariate |  |  |
| --- | --- | --- | --- | --- | --- | --- | --- |
|  | N = 485 (%) | HR | 95% CI | p-value | HR | 95% CI | p-value |
| Gender | 485 |  |  |  |  |  |  |
| female | 159 (33%) | — | — |  | — | — |  |
| male | 326 (67%) | 1.52 | 1.18,<br>1.95 | **0.001** | 1.30 | 1.00,<br>1.69 | **0.05** |
| IGHV status | 476 |  |  |  |  |  |  |
| mutated | 260 (55%) | — | — |  | — | — |  |
| unmutated | 216 (45%) | 3.33 | 2.61,<br>4.25 | **<0.001** | 2.82 | 2.15,<br>3.68 | **<0.001** |
| Age (at sample) | 485 |  |  |  |  |  |  |
| <60 | 183 (38%) | — | — |  | — | — |  |
| ≥60 | 302 (62%) | 1.10 | 0.87,<br>1.38 | 0.4 | 1.20 | 0.94,<br>1.53 | 0.14 |
| ATM status (deletion<br>and/or mutation) | 485 |  |  |  |  |  |  |
| no | 387 (80%) | — | — |  | — | — |  |
| yes | 98 (20%) | 2.08 | 1.61,<br>2.69 | **<0.001** | 1.28 | 0.97,<br>1.69 | 0.09 |
| Trisomy 12 | 485 |  |  |  |  |  |  |
| no | 417 (86%) | — | — |  | — | — |  |
| yes | 68 (14%) | 1.46 | 1.08,<br>1.97 | **0.02** | 1.40 | 1.01,<br>1.94 | **0.04** |
| Gain 8q | 485 |  |  |  |  |  |  |
| no | 477 (98%) | — | — |  |  |  |  |
| yes | 8 (1.6%) | 3.21 | 1.58,<br>6.51 | **0.001** |  |  |  |
| NOTCH1 mutation | 485 |  |  |  |  |  |  |
| no | 443 (91%) | — | — |  | — | — |  |
| yes | 42 (8.7%) | 1.98 | 1.39,<br>2.81 | **<0.001** | 1.09 | 0.75,<br>1.57 | 0.7 |
| SF3B1 mutation | 485 |  |  |  |  |  |  |
| no | 402 (83%) | — | — |  | — | — |  |
| yes | 83 (17%) | 1.95 | 1.50,<br>2.55 | **<0.001** | 1.69 | 1.28,<br>2.24 | **<0.001** |
| BIRC3 mutation | 485 |  |  |  |  |  |  |
| no | 468 (96%) | — | — |  |  |  |  |
| yes | 17 (3.5%) | 1.96 | 1.14,<br>3.35 | **0.01** |  |  |  |
| MYC expression level | 485 |  |  |  |  |  |  |
| low | 323 (67%) | — | — |  | — | — |  |
| high | 162 (33%) | 1.38 | 1.10,<br>1.74 | **0.006** | 1.28 | 1.00,<br>1.63 | **0.05** |

Abbreviations: CI = Confidence Interval, HR = Hazard Ratio

### *MYC* overexpression was associated with higher cell proliferation in *TP53*^WT^ CLL cells

To explore the biological consequences of *MYC* overexpression (*MYC*^OE^) with and without *TP53* alterations, we selected MEC-1 cell line that harbors *TP53* deletion/mutation (*TP53*^del/mut^) and PGA-1 cell line that has *TP53* wild-type (*TP53^WT^*) (Figure S2A). Initially, we successfully introduced *MYC*^OE^ in MEC-1 by CRISPR/SAM with five different sgRNAs (19, 173, 178, 240, and 369) (Figure S2B-D). The most efficient sgRNAs 19, 240, and 369 were chosen to overexpress *MYC^OE^* in both MEC-1 and PGA-1 cell lines confirmed by WB and qPCR (Figure 2A-B). In parallel, we overexpressed *MYC* by cDNA (*MYC^cOE^*) and silenced *TP53* by shRNA (*TP53*^KD^) in PGA-1 to evaluate and validate the effects of these alterations (Figure S2E).

**Figure 2:**
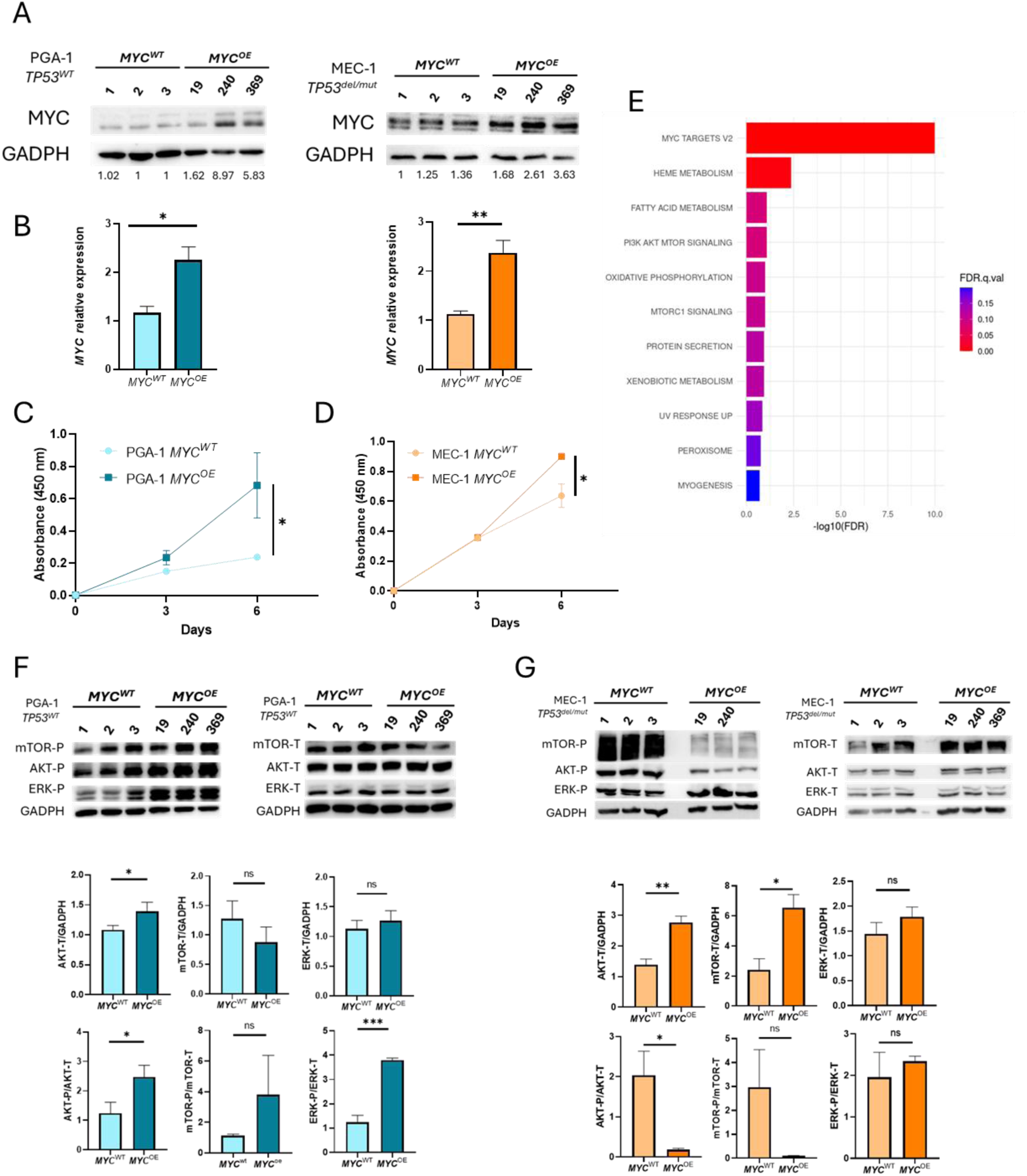
*MYC^OE^* associated with higher proliferation and AKT/mTOR activation in *TP53^WT^* background. Western Blot (A) and RTqPCR (B) of CRISPR/SAM-edited *MYC^OE^* PGA-1 and MEC-1 cell lines. Cell viability assay of CRISPR/SAM-edited MYC^OE^ PGA-1 (C) and MEC-1 (D). (E) Gene Set Enrichment Analysis of CRISPR/SAM-edited PGA-1 showing the deregulated pathways in *MYC^OE^* vs *MYC^WT^* (FDR<0.25 and p.adj<0.05). Western Blot analysis of phosphorylated and total mTOR/AKT/ERK proteins in CRISPR/SAM-edited PGA-1 (F) and MEC-1 (G) models.

Interestingly, *MYC*^OE^ induced a higher proliferation rate in the PGA-1 *TP53*^WT^ than in MEC-1 *TP53*^del/mut^ (Figure 2C-D). These results were validated in the cDNA/shRNA PGA-1 model in which we observed a significant increase of proliferation in PGA-1 *MYC*^cOE^ within the context of *TP53^WT^* (Figure S3A). These differences were not related to apoptotic defects (Figure S3B), but they could be associated with a higher proportion of cells in the S and G2/M phases in PGA-1 *TP53^WT^*-*MYC^cOE^* vs. PGA-1 *TP53^WT^*-*MYC^WT^*(Figure S3C). This difference could be related to a higher expression of the cyclin CCNE1 (Figure S3D).

### *MYC* overexpression induced transcriptomic changes related to AKT/mTOR activation in *TP53^WT^* CLL cells

Transcriptomic analysis showed that *MYC*^OE^ introduced distinct transcriptional profiles in both PGA-1 and MEC-1 models (Figure S4A-B). *MYC* overexpression by CRISPR/SAM and *TP53* status in PGA-1 and MEC1-cell line was reconfirmed by the RNA-seq results (Figure S4C). Notably, *MYC*^OE^ induced more transcriptomic changes in PGA-1 *TP53*^WT^ cellular model than in MEC-1 *TP53*^del/mut^. Whereas 82 genes were significantly deregulated in PGA-1 *TP53*^WT^-*MYC*^OE^ vs in PGA-1 *TP53*^WT^-*MYC*^WT^, no genes were significantly deregulated by *MYC*^OE^ in MEC-1 *TP53*^del/mut^ (Table S6-7). Furthermore, we discovered that *MYC* overexpression had a notable effect on gene sets associated with mTOR and PI3K/AKT pathways in the context of *TP53^WT^* (PGA-1) (Figure 2E, Table S8).

Focusing on the deregulated pathways, we validated by western-blot a considerable increased activation of AKT, mTOR and ERK when *MYC* is overexpressed in PGA-1 *TP53*^WT^ (Figure 2F). However, *MYC*^OE^ let to an inactivation of AKT/mTOR pathway as well as no differences in ERK pathway in MEC-1 *TP53*^del/mut^, as the transcriptomic data suggested (Figure 2G).

### *MYC* overexpression was related to FOXO6 upregulation in both TP53^WT^ cell lines and patients

When we compared the significant deregulated genes, identified by RNA-seq, when MYC is overexpressed in the cell line PGA-1 *TP53^WT^* and in *TP53^WT^* CLL patients, we observed an upregulation of three common genes: *FOXO6, PDE4D,* and *ARCHGAP32* (Figure 3A-B, Table S9). In addition, the expression of each gene showed a positive correlation with *MYC* expression in *TP53^WT^* CLL patients (Figure 3C and Figure S5A). Additionally, we confirmed the overexpression of these three common genes in CRISPR/SAM models by qPCR (Figure 3D and Figure S5B). Notably, *FOXO6* upregulation, which has been related to AKT/mTOR activation and cell proliferation (44), was also validated in PGA-1 *TP53^WT^-MYC^cOE^*(Figure 3E). We also confirmed this result at protein level by WB (Figure 3F). Interestingly, high levels of *FOXO6* expression were also associated with poor prognosis in TFT in *TP53^WT^* CLL patients (Figure 3G).

**Figure 3:**
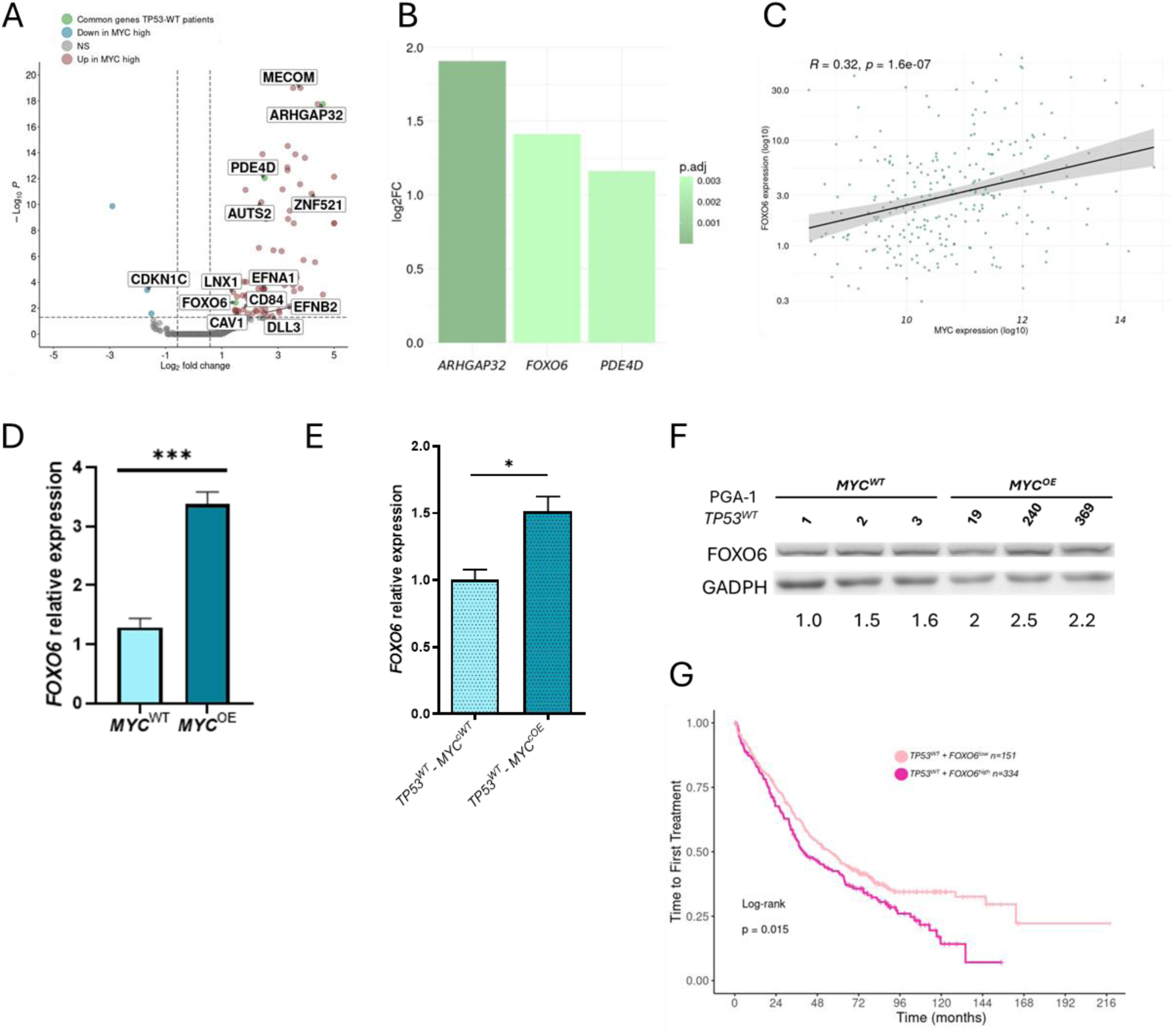
Deregulated genes in *TP53^WT^* CLL patients and cellular models. (A) Volcano plot of differential expression genes in *MYC^OE^* vs *MYC^WT^* PGA-1 cell line, showing a total of 79 significantly upregulated and 3 downregulated genes (p.adj < 0.05, absolute log₂ fold change > 0.58). Interesting candidate genes are labeled; green spots represent genes upregulated also in TP53^WT^ patients. (B) Graphic representation of log2FC values of DEGs in *TP53^WT^* CLL patients that are commonly upregulated in PGA-1 cell line. (C) Correlation analysis between *MYC* and *FOXO6* mRNA normalized expression in *TP53^WT^* CLL patients. Spearman’s correlation was calculated in patients with detected expression in both genes using Log10 values for graphs visualization. (D-E) *FOXO6* relative expression by RT-qPCR in CRISPR/SAM-edited PGA-1 (D) and shRNA/cDNA PGA-1 (E) models. (F) Western blot analysis of FOXO6 protein in CRISPR/SAM-edited PGA-1 model. (G) Kaplan–Meier plot showing the association between *FOXO6* expression and the time to first treatment in TP53^WT^ untreated CLL patients from the CLL-map portal. CLL patients were grouped according to *FOXO6* expression levels using maxstat method: *FOXO6^low^* (light pink line) and *MYC^high^* (dark pink line).

Other genes involved in cell proliferation, such as *ZNF521*, *DLL3*, *EFNB2*, *MECOM*, *EFNA1*, *LNX1*, *CD84*, *CAV1*, *ERBB4*, were also significantly upregulated (Figure 3A) and further validated in PGA-1 *TP53*^WT^-*MYC*^OE^ by qPCR (Figure S5C). In addition, *CDKN1C*, a negative regulator of proliferation, was downregulated in PGA-1 *TP53^WT^-MYC^OE^* vs *MYC^WT^* (Figure 3A, Figure S5C).

### Whereas *MYC^cOE^* cells with *TP53^WT^* background were more sensitive to BCL2i, *TP53^KD^ - MYC^cOE^* cells were resistant to BCL2i and BTKi

We assessed the impact of *MYC^OE^* on the sensitivity to current approved CLL therapies. Interestingly, we noted an increased sensitivity to venetoclax in PGA-1 *TP53^WT^*-*MYC^cOE^*compared to *TP53^WT^*-*MYC^WT^* (Figure 4A). We validated these results in CRISPR/SAM PGA-1 model (Figure S6A). Interestingly, BCL2 protein appears to be overexpressed in PGA-1 *MYC^OE^* cells (Figure 4B). Conversely, in a *TP53^KD^*context, the overexpression of *MYC* induced resistance to both venetoclax and ibrutinib (Figure 4A and 4C). These findings were also corroborated in the CRISPR/SAM MEC-1 model (Figure S6B-C). Additionally, PGA-1 *TP53^KD^/MYC^cOE^* did not have sensitivity to other BTKi (acalabrutinib, zanabrutinib, pirtobrutinib) confirming that *MYC*^OE^ in the context of *TP53* dysfunction was associated with resistance to these inhibitors (FigureS6D).

**Figure 4:**
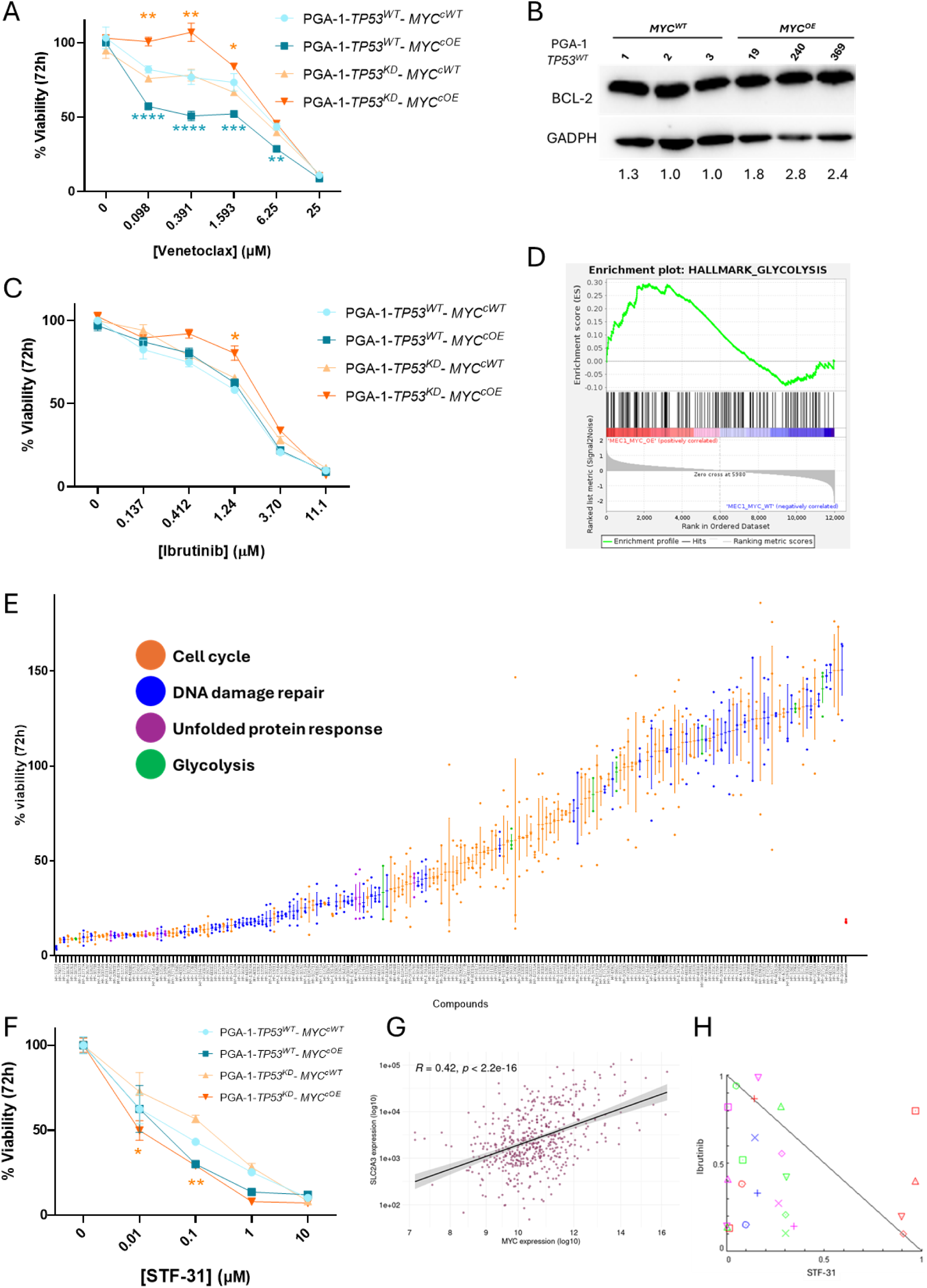
Impact of *MYC* overexpression according to *TP53* status in drug sensitivity. (A) Dose–response curves of venetoclax using increasing doses (0.098–25 µM) for 72 h in shRNA/cDNA PGA-1 models. (B) Western blot of BCL-2 in CRISPR/SAM-edited PGA-1 model. (C) Dose-response curves of ibrutinib using increasing doses (0.137–11 µM) for 72 h in shRNA/cDNA PGA-1 models. (D) Enriched glycolysis gene set from GSEA analysis in CRISPR/SAM-edited *MYC^OE^* MEC-1. (E) % cell viability of 203 drug library at 20µM in *TP53^KD^/MYC^cOE^* PGA-1. Drugs are grouped by colors in relation of the pathway in which is involved the molecular target: cell cycle (orange), DNA damage repair (blue), unfolded protein response (purple), glycolysis (green). (F) Dose-response curves of the glycolysis inhibitor STF-31 using increasing doses (0.01-10 µM) for 72 h in shRNA/cDNA PGA-1 models. (G) Correlation analysis between *MYC* and *SLC2A3* mRNA expression normalized in CLL patients. Spearman’s correlation was calculated in patients with detected expression in both genes using Log10 values for graphs visualization. (H) Normalized isobolograms for ibrutinib in combination with STF-31 in *TP53^KD^/MYC^cOE^* PGA-1 model.

### CLL cells with *MYC*^OE^ were sensitive to glycolysis inhibition independently to *TP53* status

As MYC is commonly regarded as an undruggable target, in order to find efficient therapies for CLL with the concurrence of *MYC* and *TP53* alterations, we first tested JQ1 (a BET protein inhibitor that induces cell death through *MYC* inhibition) (Figure S6E). Interestingly, PGA-1 *TP53^WT^*-*MYC^cOE^*showed higher sensitivity to this drug which could be related to MECOM overexpression in PGA-1 *TP53^WT^*-*MYC*^OE^ (Figure S5C and S6F-G; Table S6). However, PGA-1 *TP53^KD^*-*MYC^cOE^*model exhibited higher resistance to JQ1 compared to PGA-1 *TP53^KD^*-*MYC^WT^*(Figure S6E). Thus, we performed a high-throughput screening of 203 compounds targeting enriched biological processes in MEC1-*MYC^OE^* (Table S4 and S11, Figure 4D and S6H). We treated PGA-1-*TP53^KD^*-*MYC^cOE^* with drugs targeting DNA repair, unfolded protein response, glycolysis, and cell cycle (Figure 4E). The most promising treatments based on sensitivity and bibliography (abemaciclib, talazoparib, belinostat, STF-31) were tested across the four models with different alterations at increasing doses. PGA-1 *TP53^KD^*-*MYC^cOE^*model displayed less sensitivity to abemaciclib, talazoparib, and belinostat than PGA-1-*TP53^KD^*-*MYC^WT^* (Figure S6I). Remarkably, the glycolysis inhibitor STF-31, which targets SLC2A1 (GLUT1), showed increased sensitivity in PGA-1-*MYC^cOE^* independently of *TP53* status, being significant within the *TP53^KD^* context (Figure 4F). Interestingly, we observed that *MYC* expression positively correlated with the glucose transporter *SLC2A3* (*GLUT3*) in CLL patients (Figure 4G). In fact, this gene is a significantly deregulated gene in both *TP53^WT^* and *TP53^altered^* groups of CLL patients when *MYC* is overexpressed (Table S9-S10).

Subsequently, we evaluated the effect of combining STF-31 with CLL-approved drugs such as venetoclax and ibrutinib in PGA-1 *TP53^KD^-MYC^cOE^*. Whereas the combination with venetoclax showed a modest increase of the efficacy (Figure S6J), STF-31 significantly enhanced the effects of ibrutinib, indicating a synergistic interaction among both drugs (Figure 4H and S6K).

## Discussion

These results underscore the significant importance of *MYC* expression level as a prognostic indicator in CLL as well as the biological impact of its overexpression in CLL cells. Previous studies have demonstrated that 8q aberrations involving *MYC* amplification are associated with poor prognosis (15,16) and with *TP53* dysfunction in CLL (45). Furthermore, MYC activation -either by *MYC*-related chromosomal alterations or mutations in other genes such as *MGA* or *NOTCH1-* has been involved in the transformation from CLL to Richter Syndrome (20,46,47).

Building on this evidence, in this study, we further refine the prognostic role of *MYC* according to *TP53* status. Specifically, we demonstrate that CLL patients with *TP53*^WT^ who exhibit *MYC* overexpression experience significantly shorter TFT and OS, demonstrating a markedly aggressive disease course comparable to patients with *TP53* alterations. Conversely, high *MYC* levels do not appear to influence outcomes when *TP53* is altered. Taken together, these findings suggest that *MYC* overexpression may be a key driver of aggressive disease behavior in CLL, particularly in the absence of *TP53* alterations, highlighting its potential value for improved risk stratification.

The development of *MYC* overexpression models that recapitulate alterations observed in patients provides a valuable framework to explore their biological consequences and therapeutic vulnerabilities (48). While previous studies with CLL cellular models have predominantly relied on knockout systems (35,49–51), our work present, for the first time, overexpression-based approaches by CRISPR/SAM to upregulate *MYC* oncogene in two human CLL cell lines with different *TP53* backgrounds. Complementary approaches, including *MYC* overexpression via cDNA and *TP53* knockdown using shRNA, enabled the modeling of *MYC* activity in the context of monoallelic *TP53* alterations. Both *in vitro* models were applied to elucidate the molecular bases underlying the clinical differences observed in CLL patients, as well as to identify new therapeutic avenues for these high-risk CLL patients.

Interestingly, we observe markedly increased proliferation when *MYC* is overexpressed in CLL cells with *TP53*^WT^. This result can be in relation to molecular changes as we have detected an activation of AKT/mTOR signaling which is critical for cell growth and survival and has been associated with MYC activation in other cancers (52,53). This enhanced proliferation rate can also be related to a higher expression of the cyclin CCNE1, a key regulator of the G1/S transition, and a downregulation of CDKN1C, a well-characterized tumor suppressor and negative regulator of cell cycle progression, as it has been reported in other tumors (54–56). Other genes associated with cell proliferation are also upregulated in *TP53*^WT^-*MYC*^OE^ CLL cells (Figure 3A). Among them, it is worth mentioning that *ZNF521* and *MECOM* (EVI-1) have been previously implicated in leukemogenesis and promote B-cell proliferation (57–60), and *CAV1* has been associated with disease initiation and progression in *in vivo* CLL models (61). In addition, EFNA1 can positively regulate MYC activity via the AKT/mTOR pathway, raising the possibility of a feedback loop contributing to tumor progression (62,63), and DLL3 and LNX1, which are linked to NOTCH pathway, have been associated with tumor progression (64–67). In contrast, in a *TP53* altered context, the proliferative phenotype associated to *MYC* overexpression appears attenuated, suggesting that *TP53* disfunction is already associated with profound cell cycle deregulation in hematologic malignancies (65,68,69), which could limit the effects by MYC activation. Additionally, we do not detect significant transcriptomic changes associated with AKT/mTOR pathway activation on *TP53*^altered^ models when *MYC* is overexpressed, suggesting that these cells could rely on alternative oncogenic programs, reducing their dependency on MYC-driven proliferation.

Of note, higher expression levels of *FOXO6*, *PDE4D*, and *ARHGAP32* are also significantly associated with *MYC* overexpression in both our cellular model and CLL with *TP53*^WT^ background. Among them, Rho GTPase-activating proteins (ARHGAPs) and cAMP-specific phosphodiesterases (PDE4 family) play key roles in cancer progression across multiple solid tumors (70,71). However, the specific contribution of ARHGAP32 remains poorly defined. In addition, PDE4 inhibitors have shown promising effects in preclinical B-cell lymphoma models through the suppression of PI3K/AKT activity (72), suggesting the repurposing of these inhibitors for the treatment of B-cells tumors. Interestingly, FOXO6 belongs to the family of FOXO transcription factors which are essential regulators of B-cell differentiation (73). Recent studies have linked high FOXO expression to poor prognosis and tumor-promoting activity in some solid tumors and hematological malignancies, including a significant role in lymphomagenesis (74). Its overexpression can be linked to AKT/mTOR activation, since it has been demonstrated that its knockdown inhibits the activation of PI3K/Akt/mTOR pathway in colorectal cancer (44). Interestingly, we detect that FOXO6 overexpression is associated with shorter TFT in *TP53*^WT^ CLL patients. Similar observations have been reported in gastric cancer since high FOXO6 expression is linked to increased tumor aggressiveness and poorer prognosis (75). Collectively, these results indicate that *MYC* overexpression in CLL cells *TP53*^WT^ background is associated with a coordinated transcriptional reprogramming, characterized by enhanced proliferative capacity that may support tumor growth and unfavorable clinical outcomes in CLL patients harboring these alterations.

Our results also highlight the role of *MYC* expression as a predictive treatment biomarker. It has been reported that *MYC* overexpression increases sensitivity to the BCL2 inhibitor venetoclax only in the *TP53*^WT^ context that can be linked to BCL2 overexpression (76,77). These results support that CLL patients exhibiting *MYC* overexpression alongside a *TP53*^WT^ background could benefit from BCL2 inhibitor-based treatments reaching better responses than with BKT inhibitors. On the contrary, when *MYC* is overexpressed in CLL cells with *TP53* dysfunction, we observe drug resistance to both BCL2 and BTK inhibitors, highlighting that the concurrence of MYC and *TP53* alterations can significantly diminish the efficacy of current approved therapies in CLL. Our results suggest the evaluation of not only *TP53* status but also *MYC* can help on treatment decision and improve patient outcomes in CLL.

To find an effective treatment for CLLs with *MYC*^OE^ and *TP53*^altered^, we assess drug response assays with the BET inhibitor JQ1 which attenuates *MYC* expression (78,79) as well as with a library of drugs targeting the enriched pathways in MEC-1 *MYC*^OE^. Surprisingly, we only note a slight sensitivity to JQ1 when *MYC* is overexpressed in *TP53*^WT^, which can be associated with MECOM upregulation (80,81), whereas the effectiveness of this drug remains limited in *TP53*^altered^ cells. This result highlights the necessity of characterizing both *MYC* and *TP53* status to better identify drug sensitivity in CLL. Furthermore, we successfully identify the glycolysis inhibitor STF-31 as a promising approach to treat CLLs with *MYC*^OE^ independently of *TP53* status. These results can be linked to the positive correlation of *MYC* expression and the glucose transporter SLC2A3 expression observed in CLL patients. Targeting metabolism has emerged as a promising therapeutic approach focused on disrupting the energy pathways such as glycolysis (82,83). In addition, *MYC* oncogene, which can induce the expression of glucose transporters (84,85), is a key regulator of metabolism reprogramming in CLL (82). Interestingly, this inhibitor STF-31 can enhance antitumoral effects of the current CLL-approved drugs, demonstrating synergistic interactions with ibrutinib (86).

In conclusion, our results provide evidence of the clinical and biological significance of *MYC* overexpression in CLL, being more prominent in a *TP53^WT^* background. The poor prognosis associated with *MYC*^OE^ in *TP53^WT^* CLL can be linked to enhanced cell proliferation by the activation of AKT/mTOR pathway related with *FOXO6* upregulation. Regarding the predictive value of *MYC*, BCL2 inhibitors appear to be an effective choice for CLL patients exhibiting *MYC* overexpression within a *TP53^WT^* framework. Nevertheless, when *TP53* is altered, *MYC* overexpression is associated with reduced sensitivity to either BCL2 or BTK inhibitors. Additionally, we propose glycolysis inhibitors in monotherapy or combined with BTK inhibitors as a promising treatment option for CLL patients with the concurrence of *MYC* and *TP53* alterations. Thus, this study contributes to a better understanding of the heterogenous behavior of CLL, to refine the prognostic stratification based on gene expression level and to help with treatment decision based on molecular biomarkers.

## Acknowledgments

This work was supported by PID2023-149241OA-I00 and PID2024-155654OB-I00 funded by MICIU/AEI/10.13039/501100011033 by ERDF/EU, the CRIS Contra el Cáncer Foundation (2021/0088 and 2024/0031) and PIPF-CAM (PIPF-2023/SAL-GL-31039) grant from the Autonomic Community of Madrid awarded to A. Garrote-de-Barros. J. Pérez-Fernández was funded by PID2023-149241OA-I00 funded by MICIU/AEI/10.13039/501100011033 and by ERDF/EU. R. Garcia-Vicente held a Formación de Profesorado Universitario (FPU19/04933) grant from the Ministry of Science, Innovation and Universities of Spain and a grant from the Fundación Española de Hematología y Hemoterapia (FEHH). We are grateful to the Genomic Unit of the Spanish National Cancer Research Center (CNIO) for RNA sequencing. We thank Sara Izquierdo-Bermejo for technical support.

## Authors contributions

**A. Garrote-de-Barros:** Conceptualization, formal analysis, investigation, visualization, methodology, writing–original draft, writing–review and editing. **J. Pérez-Fernández:** Investigation and analysis. **A. Arroyo-Barea:** Data curation, software, formal analysis, validation, visualization. **I. Bragado-García:** Investigation. **R. Garc**í**a-Vicente:** Investigation **R. Ancos-Pintado:** Investigation. **M. Velasco-Estévez:** Investigation. **M. Linares:** Conceptualization, supervision, funding acquisition, writing–original draft, project administration, writing–review and editing. **J. Mart**í**nez-L**ó**pez:** Conceptualization, supervision, funding acquisition, writing–original draft, project administration, writing–review and editing. **M. Hernández-Sánchez:** Conceptualization, supervision, funding acquisition, writing–original draft, project administration, writing–review and editing.

## References

1. Hallek M. Chronic Lymphocytic Leukemia: 2025 Update on the Epidemiology, Pathogenesis, Diagnosis, and Therapy. Am J Hematol. 2025 Mar;100(3):450–80. doi:10.1002/ajh.27546 PubMed PMID: 39871707; PubMed Central PMCID: PMC11803567.

2. Zhu L, Fan L, Peng S, Luo X, Wen J. The Global Burden of Chronic Lymphocytic Leukemia and Its Attributable Factors in 204 Countries and Territories: Findings from the Global Burden of Disease 2021 Study and Projections to 2035 [Internet]. Research Square; 2025 [cited 2026 Feb 24]. Available from: https://www.researchsquare.com/article/rs-6751105/v1 doi:10.21203/rs.3.rs-6751105/v1

3. Braish J, Cerchione C, Ferrajoli A. An overview of prognostic markers in patients with CLL. Front Oncol. 2024 May 16;14. doi:10.3389/fonc.2024.1371057

4. Moia R, Gaidano G. Prognostication in chronic lymphocytic leukemia. Seminars in Hematology. 2024 Apr 1;Chronic Lymphocytic Leukemia (CLL)61(2):83–90. doi:10.1053/j.seminhematol.2024.02.002

5. Al-Sawaf O, Stumpf J, Zhang C, Simon F, Bosch F, Feyzi E, et al. Fixed-Duration versus Continuous Treatment for Chronic Lymphocytic Leukemia. N Engl J Med. 2026 Mar 12;394(11):1084–96. doi:10.1056/NEJMoa2515458

6. Shadman M. Diagnosis and Treatment of Chronic Lymphocytic Leukemia: A Review. JAMA. 2023 Mar 21;329(11):918–32. doi:10.1001/jama.2023.1946

7. Wood RK, Elsharawi I, Goudie M, Bruyère H, Rahmani M, Gillan TL. The ABCs of IGHV Testing in Chronic Lymphocytic Leukaemia: Current Recommendations, Ongoing Challenges, and Future Directions. International Journal of Laboratory Hematology. n/a(n/a). doi:10.1111/ijlh.70010

8. Griffin R, Wiedmeier-Nutor JE, Parikh SA, McCabe CE, O’Brien DR, Boddicker NJ, et al. Differential prognosis of single and multiple TP53 abnormalities in high-count MBL and untreated CLL. Blood Adv. 2023 Jul 3;7(13):3169–79. doi:10.1182/bloodadvances.2022009040

9. Zenz T, Eichhorst B, Busch R, Denzel T, Häbe S, Winkler D, et al. TP53 mutation and survival in chronic lymphocytic leukemia. J Clin Oncol. 2010 Oct 10;28(29):4473–9. doi:10.1200/JCO.2009.27.8762 PubMed PMID: 20697090.

10. Malcikova J, Pavlova S, Baliakas P, Chatzikonstantinou T, Tausch E, Catherwood M, et al. ERIC recommendations for TP53 mutation analysis in chronic lymphocytic leukemia—2024 update. Leukemia. 2024 Jul;38(7):1455–68. doi:10.1038/s41375-024-02267-x

11. Knisbacher BA, Lin Z, Hahn CK, Nadeu F, Duran-Ferrer M, Stevenson KE, et al. Molecular map of chronic lymphocytic leukemia and its impact on outcome. Nat Genet. 2022 Nov;54(11):1664–74. doi:10.1038/s41588-022-01140-w

12. Kittai AS, Marchetti M, Al-Sawaf O, Benjamini O, Danilov AV, Davids MS, et al. International consensus statement on diagnosis, evaluation, and research of Richter transformation: the ERIC recommendations. Blood. 2025 Jul 17;146(3):291–303. doi:10.1182/blood.2024028064

13. Pepe S, Vitale C, Giannarelli D, Visentin A, Sanna A, Frustaci AM, et al. Richter transformation in diffuse large B-cell lymphoma in patients with chronic lymphocytic leukemia receiving ibrutinib: risk factors and outcomes. Leukemia. 2025 Aug;39(8):1883–91. doi:10.1038/s41375-025-02666-8

14. Guo D, Li P, Wang Z, Li B, Bi G, Li C. MYC overexpression predicts poor prognosis and correlates with immune infiltration in chronic lymphocytic leukemia [Internet]. Research Square; 2022 [cited 2026 Feb 24]. Available from: https://www.researchsquare.com/article/rs-1276445/v1 doi:10.21203/rs.3.rs-1276445/v1

15. Ondroušková E, Bohúnová M, Závacká K, Čech P, Šmuhařová P, Boudný M, et al. Duplication of 8q24 in Chronic Lymphocytic Leukemia: Cytogenetic and Molecular Biologic Analysis of MYC Aberrations. Front Oncol. 2022 Jun 24;12. doi:10.3389/fonc.2022.859618

16. Öztürk S, Paul Y, Afzal S, Gil-Farina I, Jauch A, Bruch PM, et al. Longitudinal analyses of CLL in mice identify leukemia-related clonal changes including a Myc gain predicting poor outcome in patients. Leukemia. 2022 Feb;36(2):464–75. doi:10.1038/s41375-021-01381-4

17. Brown JR, Hanna M, Tesar B, Werner L, Pochet N, Asara JM, et al. Integrative Genomic Analysis Implicates Gain of PIK3CA at 3q26 and MYC at 8q24 in Chronic Lymphocytic Leukemia. Clin Cancer Res. 2012 Jul 15;18(14):3791–802. doi:10.1158/1078-0432.CCR-11-2342

18. Blanco G, Puiggros A, Baliakas P, Athanasiadou A, GarcÃ-a-Malo M, Collado R, et al. Karyotypic complexity rather than chromosome 8 abnormalities aggravates the outcome of chronic lymphocytic leukemia patients with TP53 aberrations. Oncotarget. 2016 Nov 4;7(49):80916–24. doi:10.18632/oncotarget.13106

19. Nguyen-Khac F. “Double-Hit” Chronic Lymphocytic Leukemia, Involving the TP53 and MYC Genes. Front Oncol. 2022 Jan 13;11. doi:10.3389/fonc.2021.826245

20. ten Hacken E, Sewastianik T, Yin S, Hoffmann GB, Gruber M, Clement K, et al. In Vivo Modeling of CLL Transformation to Richter Syndrome Reveals Convergent Evolutionary Paths and Therapeutic Vulnerabilities. Blood Cancer Discov. 2023 Mar 1;4(2):150–69. doi:10.1158/2643-3230.BCD-22-0082

21. Tsagiopoulou M, Rashmi S, Chatziaslani M, Gut I. MYC target gene activation in chronic lymphocytic leukemia and richter transformation: links to aggressiveness and tumor microenvironment interactions. Front Pharmacol. 2025 Aug 15;16. doi:10.3389/fphar.2025.1642458

22. Pozzo F, Bittolo T, Vendramini E, Bomben R, Bulian P, Rossi FM, et al. NOTCH1-mutated chronic lymphocytic leukemia cells are characterized by a MYC-related overexpression of nucleophosmin 1 and ribosome-associated components. Leukemia. 2017 Nov;31(11):2407–15. doi:10.1038/leu.2017.90

23. Close V, Close W, Kugler SJ, Reichenzeller M, Yosifov DY, Bloehdorn J, et al. FBXW7 mutations reduce binding of NOTCH1, leading to cleaved NOTCH1 accumulation and target gene activation in CLL. Blood. 2019 Feb 21;133(8):830–9. doi:10.1182/blood-2018-09-874529

24. Ritchie ME, Phipson B, Wu D, Hu Y, Law CW, Shi W, et al. limma powers differential expression analyses for RNA-sequencing and microarray studies. Nucleic Acids Res. 2015 Apr 20;43(7):e47. doi:10.1093/nar/gkv007 PubMed PMID: 25605792; PubMed Central PMCID: PMC4402510.

25. Hothorn T. maxstat: Maximally Selected Rank Statistics [Internet]. 2025 [cited 2026 May 1]. Available from: https://cran.r-project.org/web/packages/maxstat/index.html

26. Hothorn T, Zeileis A. Generalized Maximally Selected Statistics. Biometrics. 2008 Dec 1;64(4):1263–9. doi:10.1111/j.1541-0420.2008.00995.x

27. Love MI, Huber W, Anders S. Moderated estimation of fold change and dispersion for RNA-seq data with DESeq2. Genome Biol. 2014 Dec 5;15(12):550. doi:10.1186/s13059-014-0550-8

28. Robinson MD, McCarthy DJ, Smyth GK. edgeR: a Bioconductor package for differential expression analysis of digital gene expression data. Bioinformatics. 2010 Jan 1;26(1):139–40. doi:10.1093/bioinformatics/btp616

29. Stephens M. False discovery rates: a new deal. Biostatistics. 2017 Apr 1;18(2):275–94. doi:10.1093/biostatistics/kxw041

30. Doench JG, Fusi N, Sullender M, Hegde M, Vaimberg EW, Donovan KF, et al. Optimized sgRNA design to maximize activity and minimize off-target effects of CRISPR-Cas9. Nat Biotechnol. 2016 Feb;34(2):184–91. doi:10.1038/nbt.3437

31. Sanson KR, Hanna RE, Hegde M, Donovan KF, Strand C, Sullender ME, et al. Optimized libraries for CRISPR-Cas9 genetic screens with multiple modalities. Nat Commun. 2018 Dec 21;9(1):5416. doi:10.1038/s41467-018-07901-8

32. Weltner J, Balboa D, Katayama S, Bespalov M, Krjutškov K, Jouhilahti EM, et al. Human pluripotent reprogramming with CRISPR activators. Nat Commun. 2018 Jul 6;9(1):2643. doi:10.1038/s41467-018-05067-x

33. Aguilar-Garrido P, Velasco-Estévez M, Navarro-Aguadero MÁ, Otero-Sobrino Á, Ibáñez-Navarro M, Marugal MÁ, et al. The tumor suppressor HNRNPK induces p53-dependent nucleolar stress to drive ribosomopathies. J Clin Invest. 2025 Jun 16;135(12). doi:10.1172/JCI183697 PubMed PMID: 0.

34. Joung J, Konermann S, Gootenberg JS, Abudayyeh OO, Platt RJ, Brigham MD, et al. Genome-scale CRISPR-Cas9 knockout and transcriptional activation screening. Nat Protoc. 2017 Apr;12(4):828–63. doi:10.1038/nprot.2017.016

35. Quijada-Álamo M, Hernández-Sánchez M, Alonso-Pérez V, Rodríguez-Vicente AE, García-Tuñón I, Martín-Izquierdo M, et al. CRISPR/Cas9-generated models uncover therapeutic vulnerabilities of del(11q) CLL cells to dual BCR and PARP inhibition. Leukemia. 2020 Jun;34(6):1599–612. doi:10.1038/s41375-020-0714-3

36. Ewels P, Magnusson M, Lundin S, Käller M. MultiQC: summarize analysis results for multiple tools and samples in a single report. Bioinformatics. 2016 Oct 1;32(19):3047–8. doi:10.1093/bioinformatics/btw354

37. Wingett SW, Andrews S. FastQ Screen: A tool for multi-genome mapping and quality control. F1000Res. 2018;7:1338. doi:10.12688/f1000research.15931.2 PubMed PMID: 30254741; PubMed Central PMCID: PMC6124377.

38. Smith T, Heger A, Sudbery I. UMI-tools: modeling sequencing errors in Unique Molecular Identifiers to improve quantification accuracy. Genome Res. 2017 Mar;27(3):491–9. doi:10.1101/gr.209601.116 PubMed PMID: 28100584; PubMed Central PMCID: PMC5340976.

39. Dobin A, Davis CA, Schlesinger F, Drenkow J, Zaleski C, Jha S, et al. STAR: ultrafast universal RNA-seq aligner. Bioinformatics. 2013 Jan 1;29(1):15–21. doi:10.1093/bioinformatics/bts635

40. Liao Y, Smyth GK, Shi W. featureCounts: an efficient general purpose program for assigning sequence reads to genomic features. Bioinformatics. 2014 Apr 1;30(7):923–30. doi:10.1093/bioinformatics/btt656

41. Subramanian A, Tamayo P, Mootha VK, Mukherjee S, Ebert BL, Gillette MA, et al. Gene set enrichment analysis: A knowledge-based approach for interpreting genome-wide expression profiles. Proceedings of the National Academy of Sciences. 2005 Oct 25;102(43):15545–50. doi:10.1073/pnas.0506580102

42. Therneau TM, Grambsch PM. Modeling Survival Data: Extending the Cox Model [Internet]. New York, NY: Springer; 2000 [cited 2026 May 1]. (Dietz K, Gail M, Krickeberg K, Samet J, Tsiatis A, editors. Statistics for Biology and Health). Available from: http://link.springer.com/10.1007/978-1-4757-3294-8 doi:10.1007/978-1-4757-3294-8

43. Drawing Survival Curves using ggplot2 [Internet]. [cited 2026 May 1]. Available from: https://rpkgs.datanovia.com/survminer/index.html

44. Li Q, Tang H, Hu F, Qin C. Silencing of FOXO6 inhibits the proliferation, invasion, and glycolysis in colorectal cancer cells. Journal of Cellular Biochemistry. 2019;120(3):3853–60. doi:10.1002/jcb.27667

45. Puiggros A, Collado R, Calasanz MJ, Ortega M, Ruiz-Xivillé N, Rivas-Delgado A, et al. Patients with chronic lymphocytic leukemia and complex karyotype show an adverse outcome even in absence of TP53/ATM FISH deletions. Oncotarget. 2017 Apr 21;8(33):54297–303. doi:10.18632/oncotarget.17350 PubMed PMID: 28903342; PubMed Central PMCID: PMC5589581.

46. Iyer P, Zhang B, Liu T, Jin M, Hart K, Zhang J, et al. MGA deletion leads to Richter’s transformation by modulating mitochondrial OXPHOS. Sci Transl Med. 2024 Jul 31;16(758):eadg7915. doi:10.1126/scitranslmed.adg7915 PubMed PMID: 39083585; PubMed Central PMCID: PMC12377275.

47. Nadeu F, Efremov DG. The biology and evolution of Richter transformation in chronic lymphocytic leukemia. Seminars in Cancer Biology. 2026 Mar 1;120:48–60. doi:10.1016/j.semcancer.2026.02.002

48. Katti A, Diaz BJ, Caragine CM, Sanjana NE, Dow LE. CRISPR in cancer biology and therapy. Nat Rev Cancer. 2022 May;22(5):259–79. doi:10.1038/s41568-022-00441-w

49. Hernández-Sánchez M. CRISPR/Cas9 in Chronic Lymphocytic Leukemia. Encyclopedia. 2022 Jun;2(2):928–36. doi:10.3390/encyclopedia2020061

50. Rodríguez-Sánchez A, Quijada-Álamo M, Pérez-Carretero C, Herrero AB, Arroyo-Barea A, Dávila-Valls J, et al. SAMHD1 dysfunction impairs DNA damage response and increases sensitivity to PARP inhibition in chronic lymphocytic leukemia. Sci Rep. 2025 Mar 26;15(1):10446. doi:10.1038/s41598-025-93629-7

51. Arruga F, Gizdic B, Bologna C, Cignetto S, Buonincontri R, Serra S, et al. Mutations in NOTCH1 PEST domain orchestrate CCL19-driven homing of chronic lymphocytic leukemia cells by modulating the tumor suppressor gene DUSP22. Leukemia. 2017 Sep;31(9):1882–93. doi:10.1038/leu.2016.383

52. Liu J, Feng W, Liu M, Rao H, Li X, Teng Y, et al. Stomach-specific c-Myc overexpression drives gastric adenoma in mice through AKT/mammalian target of rapamycin signaling. Bosn J Basic Med Sci. 2021 Aug;21(4):434–46. doi:10.17305/bjbms.2020.4978 PubMed PMID: 33259779; PubMed Central PMCID: PMC8292868.

53. Kumar D, Kanchan R, Chaturvedi NK. Targeting protein synthesis pathways in MYC-amplified medulloblastoma. Discov Onc. 2025 Jan 8;16(1):23. doi:10.1007/s12672-025-01761-7

54. Zhao H, Wang J, Zhang Y, Yuan M, Yang S, Li L, et al. Prognostic Values of *CCNE1* Amplification and Overexpression in Cancer Patients: A Systematic Review and Meta-analysis. Journal of Cancer. 2018 Jun 14;9(13):2397–407. doi:10.7150/jca.24179

55. Gorski JW, Ueland FR, Kolesar JM. CCNE1 Amplification as a Predictive Biomarker of Chemotherapy Resistance in Epithelial Ovarian Cancer. Diagnostics. 2020 May;10(5):279. doi:10.3390/diagnostics10050279

56. Stampone E, Caldarelli I, Zullo A, Bencivenga D, Mancini FP, Della Ragione F, et al. Genetic and Epigenetic Control of CDKN1C Expression: Importance in Cell Commitment and Differentiation, Tissue Homeostasis and Human Diseases. International Journal of Molecular Sciences. 2018 Apr;19(4):1055. doi:10.3390/ijms19041055

57. Qin R, Yang T, Jiang H, Yu M. ZNF521 promotes acute myeloid leukemogenesis by suppressing the expression and acetylation of SMC3. Heliyon. 2024 Sep 5;10(18):e37528. doi:10.1016/j.heliyon.2024.e37528 PubMed PMID: 39309877; PubMed Central PMCID: PMC11415694.

58. Al Dallal S, Wolton K, Hentges KE. *Zfp521* promotes B-cell viability and *cyclin D1* gene expression in a B cell culture system. Leukemia Research. 2016 Jul 1;46:10–7. doi:10.1016/j.leukres.2016.03.013

59. Schmoellerl J, Barbosa IAM, Minnich M, Andersch F, Smeenk L, Havermans M, et al. EVI1 drives leukemogenesis through aberrant ERG activation. Blood. 2023 Feb 2;141(5):453–66. doi:10.1182/blood.2022016592

60. Masamoto Y, Chiba A, Mizuno H, Hino T, Hayashida H, Sato T, et al. EVI1 exerts distinct roles in AML via ERG and cyclin D1 promoting a chemoresistant and immune-suppressive environment. Blood Adv. 2023 Apr 14;7(8):1577–93. doi:10.1182/bloodadvances.2022008018

61. Shukla A, Cutucache CE, Sutton GL, Pitner MA, Rai K, Rai S, et al. Absence of caveolin-1 leads to delayed development of chronic lymphocytic leukemia in Eμ-TCL1 mouse model. Experimental Hematology. 2016 Jan 1;44(1):30–37.e1. doi:10.1016/j.exphem.2015.09.005

62. Jiang H, Wang S, Liu Y, Zheng C, Chen L, Zheng K, et al. Targeting EFNA1 suppresses tumor progression via the cMYC-modulated cell cycle and autophagy in esophageal squamous cell carcinoma. Discov Onc. 2023 May 9;14(1):64. doi:10.1007/s12672-023-00664-9

63. Ondrisova L, Seda V, Hlavac K, Pavelkova P, Hoferkova E, Chiodin G, et al. FoxO1/Rictor axis induces a nongenetic adaptation to ibrutinib via Akt activation in chronic lymphocytic leukemia. J Clin Invest. 134(23):e173770. doi:10.1172/JCI173770 PubMed PMID: 39436708; PubMed Central PMCID: PMC11601945.

64. Furuta M, Kikuchi H, Shoji T, Takashima Y, Kikuchi E, Kikuchi J, et al. DLL3 regulates the migration and invasion of small cell lung cancer by modulating Snail. Cancer Science. 2019;110(5):1599–608. doi:10.1111/cas.13997

65. Zhang Y, Shang L, Han J, Shen X, Liu H, Yang J, et al. Biological and immunological significance of DLL3 expression in different tumor tissues: a pan-cancer analysis. Aging. 2023 Apr 22;15(9):3427–41. doi:10.18632/aging.204672 PubMed PMID: 37179118.

66. Jang M, Park R, Park YI, Park Y, Lee JI, Namkoong S, et al. LNX1 Contributes to Cell Cycle Progression and Cisplatin Resistance. Cancers (Basel). 2021 Aug 12;13(16):4066. doi:10.3390/cancers13164066 PubMed PMID: 34439220; PubMed Central PMCID: PMC8394373.

67. Yuan C, Chang K, Xu C, Li Q, Du Z. High expression of DLL3 is associated with a poor prognosis and immune infiltration in invasive breast cancer patients. Transl Oncol. 2021 Apr 26;14(7):101080. doi:10.1016/j.tranon.2021.101080 PubMed PMID: 33915517; PubMed Central PMCID: PMC8093948.

68. Ahmadi SE, Rahimian E, Rahimi S, Zarandi B, Bahraini M, Soleymani M, et al. From regulation to deregulation of p53 in hematologic malignancies: implications for diagnosis, prognosis and therapy. Biomark Res. 2024 Nov 14;12(1):137. doi:10.1186/s40364-024-00676-9

69. Wang H, Guo M, Wei H, Chen Y. Targeting p53 pathways: mechanisms, structures and advances in therapy. Sig Transduct Target Ther. 2023 Mar 1;8(1):92. doi:10.1038/s41392-023-01347-1

70. Hsien Lai S, Zervoudakis G, Chou J, Gurney ME, Quesnelle KM. PDE4 subtypes in cancer. Oncogene. 2020 May;39(19):3791–802. doi:10.1038/s41388-020-1258-8

71. Yang C, Wu S, Mou Z, Zhou Q, Zhang Z, Chen Y, et al. Transcriptomic Analysis Identified ARHGAP Family as a Novel Biomarker Associated With Tumor-Promoting Immune Infiltration and Nanomechanical Characteristics in Bladder Cancer. Front Cell Dev Biol. 2021 Jul 7;9. doi:10.3389/fcell.2021.657219

72. Kelly K, Mejia A, Suhasini AN, Lin AP, Kuhn J, Karnad AB, et al. Safety and Pharmacodynamics of the PDE4 Inhibitor Roflumilast in Advanced B-cell Malignancies. Clin Cancer Res. 2017 Feb 28;23(5):1186–92. doi:10.1158/1078-0432.CCR-16-1207

73. Ushmorov A, Wirth T. FOXO in B-cell lymphopoiesis and B cell neoplasia. Seminars in Cancer Biology. 2018 Jun 1;FOXO Family in Cancer50:132–41. doi:10.1016/j.semcancer.2017.07.008

74. Kabrani E, Chu VT, Tasouri E, Sommermann T, Baßler K, Ulas T, et al. Nuclear FOXO1 promotes lymphomagenesis in germinal center B cells. Blood. 2018 Dec 20;132(25):2670–83. doi:10.1182/blood-2018-06-856203

75. Wang JH, Tang H sheng, Li XS, Zhang XL, Yang XZ, Zeng LS, et al. Elevated FOXO6 expression correlates with progression and prognosis in gastric cancer. Oncotarget. 2017 Mar 6;8(19):31682–91. doi:10.18632/oncotarget.15920 PubMed PMID: 28404958; PubMed Central PMCID: PMC5458239.

76. Rodríguez-Medina C, Stuckey R, Bilbao-Sieyro C, Gómez-Casares MT. Biomarkers of Response to Venetoclax Therapy in Acute Myeloid Leukemia. Int J Mol Sci. 2024 Jan 24;25(3):1421. doi:10.3390/ijms25031421 PubMed PMID: 38338698; PubMed Central PMCID: PMC10855565.

77. Pan R, Hogdal LJ, Benito JM, Bucci D, Han L, Borthakur G, et al. Selective BCL-2 Inhibition by ABT-199 Causes On-Target Cell Death in Acute Myeloid Leukemia. Cancer Discov. 2014 Mar 3;4(3):362–75. doi:10.1158/2159-8290.CD-13-0609

78. Delmore JE, Issa GC, Lemieux ME, Rahl PB, Shi J, Jacobs HM, et al. BET bromodomain inhibition as a therapeutic strategy to target c-Myc. Cell. 2011 Sep 16;146(6):904–17. doi:10.1016/j.cell.2011.08.017 PubMed PMID: 21889194; PubMed Central PMCID: PMC3187920.

79. Shao Q, Kannan A, Lin Z, Stack BC, Suen JY, Gao L. BET protein inhibitor JQ1 attenuates Myc-amplified MCC tumor growth in vivo. Cancer Res. 2014 Dec 1;74(23):7090–102. doi:10.1158/0008-5472.CAN-14-0305 PubMed PMID: 25277525; PubMed Central PMCID: PMC4322674.

80. Chen Y, Jiang Q, Xue Y, Chen W, Hua M. CRISPR-Cas9-mediated deletion enhancer of MECOM play a tumor suppressor role in ovarian cancer. Funct Integr Genomics. 2024 Jul 12;24(4):125. doi:10.1007/s10142-024-01399-8

81. Birdwell CE, Fiskus W, Mill CP, Kadia TM, Daver N, DiNardo CD, et al. BET inhibitor-based combinations targeting novel dependencies in MECOM-rearranged (r) AML. Leukemia. 2026 Feb;40(2):304–13. doi:10.1038/s41375-025-02842-w

82. Simon-Molas H, Montironi C, Kabanova A, Eldering E. Metabolic reprogramming in the CLL TME; potential for new therapeutic targets. Seminars in Hematology. 2024 Jun 1;Chronic Lymphocytic Leukemia (CLL) Part 261(3):155–62. doi:10.1053/j.seminhematol.2024.02.001

83. Nie Y, Yun X, Zhang Y, Wang X. Targeting metabolic reprogramming in chronic lymphocytic leukemia. Exp Hematol Oncol. 2022 Jun 27;11:39. doi:10.1186/s40164-022-00292-z PubMed PMID: 35761419; PubMed Central PMCID: PMC9235173.

84. Heydarzadeh S, Moshtaghie AA, Daneshpoor M, Hedayati M. Regulators of glucose uptake in thyroid cancer cell lines. Cell Commun Signal. 2020 Jun 3;18(1):83. doi:10.1186/s12964-020-00586-x

85. Marengo B, Garbarino O, Speciale A, Monteleone L, Traverso N, Domenicotti C. MYC Expression and Metabolic Redox Changes in Cancer Cells: A Synergy Able to Induce Chemoresistance. Oxidative Medicine and Cellular Longevity. 2019;2019(1):7346492. doi:10.1155/2019/7346492

86. Galicia-Vázquez G, Smith S, Aloyz R. Del11q-positive CLL lymphocytes exhibit altered glutamine metabolism and differential response to GLS1 and glucose metabolism inhibition. Blood Cancer Journal. 2018 Jan 24;8(1):13. doi:10.1038/s41408-017-0039-2

