## Supplementary material for "MYC overexpression drives poor prognosis and differential sensitivity to treatments according to TP53 status in chronic lymphocytic leukemia"

**Supplementary methods**

**Bioinformatic analysis from RNA-seq data of cellular models**

Raw sequencing data in FASTQ format were obtained for each sample and subjected to quality assessment using FastQC (v. 0.12.1; RRID:SCR_014583) (1). MultiQC (v.1.18; RRID:SCR_014982) (2) was used for aggregation of quality control metrics.

Sequencing reads underwent preprocessing to extract UMIs and to remove low-quality sequences, adapters, and polyA tails using UMI-tools and Bbduk (BBMap v.39.01; RID: SCR_016965) with customized parameters (k = 13, ktrim = r, mink = 5, qtrim = r, trimq = 20, minlength = 20). FastQ Screen (v0.15.3; RRID:SCR_000141) (3) was employed to assess contamination, using the Bowtie2 aligner (--aligner “bowtie2”) (RRID:SCR_016368).

Cleaned reads were aligned to the human genome assembly GRCh38 using STAR (v. 2.7.11a; RRID:SCR_004463) (4), with customized parameters (--outFilterType BySJout, --outFilterMultimapNmax 25, --alignSJoverhangMin 8, --alignSJDBoverhangMin 1, --outFilterMismatchNmax 999, --alignIntronMin 20, --alignIntronMax 1000000, --alignMatesGapMax 1000000, --outSAMattributes NH HI AS NM MD, -outSAMtype BAM SortedByCoordinate) suitable for 3’ RNA-seq (5). UMI-based deduplication was performed with UMI-tools and alignment quality was evaluated using Qualimap2 (v. 2.3; RRID:SCR_026258) (6,7) with default parameters, and with samtools stats and samtools flagstat (8) (RRID:SCR_002105) also using default parameters, and strand-specificity was assessed with RSeqQC (v. 5.0.1; RRID:SCR_026511) (9) using the parameter -p strand-specific-forward. A high-quality alignment was characterized by meeting the following criteria: a unique mapping rate exceeding 70%, a strandness designated as "SingleEnd", and a significant proportion of sequences (~90%) mapped to exons.

Gene-level quantification was performed using featureCounts (v.2.0.6; RRID:SCR_012919) (10) from the Subread package (RRID:SCR_009803) based on the ENSEMBL (RRID:SCR_002344) human GRCh38 gene annotations (release 111). Multi-mapping reads were assigned to all overlapping features (-O) with strand-specificity specified (-s 1).

Feature counts and clinical data were processed in R (v.4.3.1; RRID:SCR_001905). Prior to differential expression analysis, low expressed genes were filtered using filterByExpr from edgeR (v.4.0.16; RRID:SCR_012802) (11), which removes genes with insufficient expression in a number of samples across the experimental groups to improve the statistical power of the analysis. This filtering step retained 14315 out of 63241 genes. Exploratory analysis was conducted using PCA with plotPCA to visualize underlying factors influencing gene expression profiles.

Differential gene expression analysis was performed using DESeq2 (v.1.42.1; RRID:SCR_015687) (12), incorporating log fold-change shrinkage with ashr (13), the impact of *MYC* overexpression in DEG was evaluated individually in PGA-1 and MEC-1. Genes were considered differentially expressed with $|log2FC| > 0.58$ and adjusted $P < 0.05$.

Gene set enrichment analysis (GSEA) was performed using the GSEA Desktop application (v4.3.2; RRID:SCR_003199) (14). Analyses were conducted on ranked gene lists derived from differential expression analyses of cell line datasets. Comparisons were performed between $MYC^{OE}$ and $MYC^{WT}$ conditions in each cellular model. Enrichment was assessed using the Hallmark gene set collection (MSigDB) with 1,000 permutations. Pathways with a normalized enrichment score (NES), false discovery rate (FDR) < 0.25, and adjusted p-value < 0.05 were considered significant.

All figures related to RNA-seq analysis were generated using R software (v.4.4.3; RRID:SCR_001905; EnhancedVolcano (v.1.24.0; RRID:SCR_018931) (15), and ggplot2 (v. 3.5.1; RRID:SCR_014601) (16) packages.

#### References


1. Babraham Bioinformatics - FastQC A Quality Control tool for High Throughput Sequence Data [Internet]. [cited 2026 May 1]. Available from: https://www.bioinformatics.babraham.ac.uk/projects/fastqc/

2. Ewels P, Magnusson M, Lundin S, Käller M. MultiQC: summarize analysis results for multiple tools and samples in a single report. Bioinformatics. 2016 Oct 1;32(19):3047–8. doi:10.1093/bioinformatics/btw354

3. Wingett SW, Andrews S. FastQ Screen: A tool for multi-genome mapping and quality control. F1000Res. 2018;7:1338. doi:10.12688/f1000research.15931.2 PubMed PMID: 30254741; PubMed Central PMCID: PMC6124377.

4. Dobin A, Davis CA, Schlesinger F, Drenkow J, Zaleski C, Jha S, et al. STAR: ultrafast universal RNA-seq aligner. Bioinformatics. 2013 Jan 1;29(1):15–21. doi:10.1093/bioinformatics/bts635

5. Paropkari AD, Bapat PS, Sindi SS, Nobile CJ. A computational workflow for the analysis of 3' Tag-Seq data. Curr Protoc. 2023 Feb;3(2):e664. doi:10.1002/cpz1.664 PubMed PMID: 36779816; PubMed Central PMCID: PMC9930165.

6. Smith T, Heger A, Sudbery I. UMI-tools: modeling sequencing errors in Unique Molecular Identifiers to improve quantification accuracy. Genome Res. 2017 Mar;27(3):491–9. doi:10.1101/gr.209601.116 PubMed PMID: 28100584; PubMed Central PMCID: PMC5340976.

7. Okonechnikov K, Conesa A, García-Alcalde F. Qualimap 2: advanced multi-sample quality control for high-throughput sequencing data. Bioinformatics. 2016 Jan 15;32(2):292–4. doi:10.1093/bioinformatics/btv566 PubMed PMID: 26428292; PubMed Central PMCID: PMC4708105.

8. Danecek P, Bonfield JK, Liddle J, Marshall J, Ohan V, Pollard MO, et al. Twelve years of SAMtools and BCFtools. Gigascience. 2021 Feb 16;10(2):giab008. doi:10.1093/gigascience/giab008 PubMed PMID: 33590861; PubMed Central PMCID: PMC7931819.

9. RSeQC: quality control of RNA-seq experiments | Bioinformatics | Oxford Academic [Internet]. [cited 2026 May 1]. Available from: https://academic.oup.com/bioinformatics/article/28/16/2184/325191?login=true

10. Liao Y, Smyth GK, Shi W. featureCounts: an efficient general purpose program for assigning sequence reads to genomic features. Bioinformatics. 2014 Apr 1;30(7):923–30. doi:10.1093/bioinformatics/btt656

11. Robinson MD, McCarthy DJ, Smyth GK. edgeR: a Bioconductor package for differential expression analysis of digital gene expression data. Bioinformatics. 2010 Jan 1;26(1):139–40. doi:10.1093/bioinformatics/btp616

12. Love MI, Huber W, Anders S. Moderated estimation of fold change and dispersion for RNA-seq data with DESeq2. Genome Biol. 2014 Dec 5;15(12):550. doi:10.1186/s13059-014-0550-8

13. Stephens M. False discovery rates: a new deal. Biostatistics. 2017 Apr 1;18(2):275–94. doi:10.1093/biostatistics/kxw041

14. Subramanian A, Tamayo P, Mootha VK, Mukherjee S, Ebert BL, Gillette MA, et al. Gene set enrichment analysis: A knowledge-based approach for interpreting genome-wide expression profiles. Proceedings of the National Academy of Sciences. 2005 Oct 25;102(43):15545–50. doi:10.1073/pnas.0506580102

15. Blighe K. kevinblighe/EnhancedVolcano [R] [Internet]. 2026 [cited 2026 May 1]. Available from: https://github.com/kevinblighe/EnhancedVolcano

16. Wickham H. ggplot2 [Internet]. Cham: Springer International Publishing; 2016 [cited 2026 May 1]. (Use R!). Available from: http://link.springer.com/10.1007/978-3-319-24277-4 doi:10.1007/978-3-319-24277-4

### Supplementary Tables:

| Target | Sequence 5'->3' |
|---|---|
| MYC 19 | ATCGCGCTGAGTATAAAAGC |
| MYC 173 | CCCTTTATAATGCGAGGGTC |
| MYC 178 | ACCGGCCCTTTATAATGCGA |
| MYC 240 | GGTTCCCAAAGCAGAGGGCG |
| MYC 369 | GCGCGCGTAGTTAATTCATG |

**Supplementary Table 1:** Sequences of MYC sgRNAs for the generation by CRISPR/SAM CLL cellular models.

| Primers Name | Sequence 5' -> 3' |
|---|---|
| qPCR_hSAC3D1_Rev | CACTGCACAGCGCAACTTG |
| qPCR_hASC3D1_For | AAGCCCTGCATGAGGTTCTAC |
| qPCR_hEFNB2_For | TATGCAGAACTGCGATTTCCAA |
| qPCR_hEFNB2_Rev | TGGGTATAGTACCAGTCCTTGTC |
| qPCR_hLNX1_For | TGGTGGGAGGTAGCGAAAC |
| qPCR_hLNX1_Rev | TCCATCCCGTTGACCTTTAGAAT |
| qPCR_hCDKN1C_For | GCGGCGATCAAGAAGCTGT |
| qPCR_hCDKN1C_Rev | GCTTGGCGAAGAAATCGGAGA |
| qPCR_hDLL3_For | CACTCCCGGATGCACTCAAC |
| qPCR_hDLL3_Rev | GATTCCAATCTACGGACGAGC |
| qPCR_hERBB4_For | GCAGATGCTACGGACCTTACG |
| qPCR_hERBB4_Rev | GACACTGAGTAACACATGCTCC |
| qPCR_hCAV1_For | GCGACCCTAAACACCTCAAC |
| qPCR_hCAV1_Rev | ATGCCGTCAAAACTGTGTGTC |
| qPCR_hMECOM-For | TATCCACGAAGAACGGCAATATC |
| qPCR_hMECOM-Rev | CATGGAAACTTTTGGTGATCTGC |
| qPCR_hZNF521_For | CAACTGACAGATGGAGTGGATG |
| qPCR_hZNF521_Rev | GCTAGGGGAAGTCTGATCCTT |
| qPCR_hCD84_For | GGAGAAGAGGGTAATGTCCTTCA |
| qPCR_hCD84_Rev | CCATTGCGATGTCTGCACA |
| qPCR_hEFNA1_For | TCAGGCCCATGACAATCCAC |
| qPCR_hEFNA1_Rev | GTGACCGATGCTATGTAGAACC |
| qPCR_hFOXO6_for | TCTACGACTGGATGGTCCGT |
| qPCR_hFOXO6_rev | TGTGCCGGATGGAGTTCTTC |
| qPCR_hARHGAP32_For | TCATCCTGAATCACGTTGATGTG |
| qPCR_hARHGAP32_Rev | GGACTTGGGCCTTGATAGAGAAG |
| qPCR_hTP53_For | CAGCACATGACGGAGGTTGT |
| qPCR_hTP53_Rev | TCATCCAAATACTCCACACGC |
| qPCR_hPDE4D_For | GACCAATGTCTCAGATCAGTGG |
| qPCR_hPDE4D_Rev | GTCAAGGGCCGGTTACCAG |
| qPCR_hMYC_Fow | GGCTCCTGGCAAAAGGTCA |
| qPCR_hMYC_Rev | CTGCGTAGTTGTGCTGATGT |
| qPCR_hGAPDH_Fow | CTGGGCTACACTGAGCACC |
| qPCR_hGAPDH_Rev | AAGTGGTCGTTGAGGGCAATG |

**Supplementary Table 2**: Sequences of qPCR primers

| Antibody | Host | Clone | Dilution | Reference |
|---|---|---|---|---|
| Anti-GADPH | Rabbit | 14C10 | 1:1500 | CellSignaling #2118 |
| Anti-c-MYC | Rabbit | Y69 | 1:1000 | Abcam #AB32072 |
| Anti-ERK (p44/42 MAPK) | Rabbit | 137F5 | 1:1000 | CellSignaling #4695 |
| Anti-P-ERK (Phospho-p44/42 MAPK) - (Thr202/Tyr204) | Rabbit | D13.14.4E | 1:2000 | CellSignaling #4370 |
| Anti-mTOR | Rabbit | Polyclonal | 1:1000 | CellSignaling #2972 |
| Anti-P-mTOR - (Ser2448) | Rabbit | Polyclonal | 1:400 | CellSignaling #2971 |
| Anti-AKT | Rabbit | C67E7 | 1:1000 | CellSignaling #4691 |
| Anti-P-AKT - (Ser473) | Rabbit | D9E | 1:2000 | CellSignaling #4060 |
| Anti-BCL-2 | Rabbit | D55G8 | 1:1000 | CellSignaling #4223 |
| Anti-MECOM (Anti-EVI-1) | Rabbit | C50E12 | 1:500 | CellSignaling #2593 |
| Anti-FOXO6 | Rabbit | Polyclonal | 1:800 | Proteintech #19122-1-AP |
| Secondary Anti-rabbit | Goat | Polyclonal | 1:5000 | CellSignaling #7074 |

**Supplementary Table 3:** Antibodies used in Western Blot

| Variable | Cohort Overall | COX-TFT-Univariant | | | COX-TFT-Multivariant | | |
|---|---|---|---|---|---|---|---|
| | N = 531 (%) | HR | 95% CI | p-value | HR | 95% CI | p-value |
| Gender | 531 | | | | | | |
| female | 175 (33%) | — | — | | — | — | |
| male | 356 (67%) | 1.48 | 1.17, 1.87 | **0.001** | 1.25 | 0.98, 1.60 | 0.07 |
| IGHV status | 521 | | | | | | |
| mutated | 277 (53%) | — | — | | — | — | |
| unmutated | 244 (47%) | 3.28 | 2.61, 4.13 | **<0.001** | 2.69 | 2.09, 3.47 | **<0.001** |
| Age (at sample) | 531 | | | | | | |
| <60 | 196 (37%) | — | — | | — | — | |
| ≥60 | 335 (63%) | 1.05 | 0.85, 1.31 | 0.6 | 1.12 | 0.90, 1.41 | 0.3 |
| TP53 status (deletion and/or | 531 | | | | | | |
| no | 485 (91%) | — | — | | — | — | |
| yes | 46 (8.7%) | 1.88 | 1.33, 2.65 | **<0.001** | 1.55 | 1.07, 2.23 | **0.02** |
| ATM status (deletion and/or mutation) | 531 | | | | | | |
| no | 425 (80%) | — | — | | — | — | |
| yes | 106 (20%) | 2.05 | 1.60, 2.61 | **<0.001** | 1.31 | 1.00, 1.70 | **0.05** |
| Trisomy 12 | 531 | | | | | | |
| no | 461 (87%) | — | — | | — | — | |
| yes | 70 (13%) | 1.40 | 1.05, 1.88 | **0.02** | 1.49 | 1.08, 2.05 | **0.01** |
| Gain 8q | 531 | | | | | | |
| no | 519 (98%) | — | — | | | | |
| yes | 12 (2.3%) | 2.32 | 1.27, 4.23 | **0.006** | | | |
| NOTCH1 mutation | 531 | | | | | | |
| no | 484 (91%) | — | — | | — | — | |
| yes | 47 (8.9%) | 1.92 | 1.38, 2.67 | **<0.001** | 1.14 | 0.81, 1.60 | 0.5 |
| SF3B1 mutation | 531 | | | | | | |
| no | 437 (82%) | — | — | | — | — | |
| yes | 94 (18%) | 1.98 | 1.54, 2.54 | **<0.001** | 1.71 | 1.31, 2.22 | **<0.001** |
| BIRC3 mutation | 531 | | | | | | |
| no | 513 (97%) | — | — | | | | |
| yes | 18 (3.4%) | 1.91 | 1.14, 3.22 | **0.01** | | | |
| MYC expression level | 531 | | | | | | |
| low | 343 (65%) | — | — | | — | — | |
| high | 188 (35%) | 1.32 | 1.06, 1.64 | **0.01** | 1.17 | 0.92, 1.48 | 0.2 |

Abbreviations: CI = Confidence Interval, HR = Hazard Ratio

**Supplementary Table 5:** COX model for Time to First Treatment of 531 untreated CLL patients from CLL-map portal

| Pathway Name | NES | SIZE | FDR.q.val | NOM.p.val |
|---|---|---|---|---|
| MYC TARGETS V2 | 2.137928 | 57 | 0 | 0 |
| MYOGENESIS | 1.4484737 | 87 | 0.19849756 | 0.027941177 |
| UV RESPONSE UP | 1.4042509 | 115 | 0.14532809 | 0.028129395 |
| XENOBIOTIC METABOLISM | -1.3546646 | 110 | 0.12036746 | 0.036666665 |
| MTORC1 SIGNALING | -1.3861656 | 193 | 0.10479097 | 0.007751938 |
| PI3K AKT MTOR SIGNALING | -1.4314656 | 91 | 0.0851145 | 0.015243903 |
| FATTY ACID METABOLISM | -1.4576938 | 114 | 0.083791986 | 0.013513514 |
| OXIDATIVE PHOSPHORYLATION | -1.4616991 | 195 | 0.100573614 | 0 |
| PROTEIN SECRETION | -1.4758887 | 86 | 0.12003101 | 0.013071896 |
| PEROXISOME | -1.4786979 | 77 | 0.17486058 | 0.011940299 |
| HEME METABOLISM | -1.875929 | 141 | 0.0042450977 | 0 |

**Supplementary Table 8:** Enriched pathways in MEC-1 MYC-OE vs MYC-WT

| Pathway Name | NES | SIZE | FDR.q.val | NOM.p.val |
|---|---|---|---|---|
| MYC TARGETS V2 | 2.150431 | 56 | 0.0006521739 | 0 |
| MYC TARGETS V1 | 2.108725 | 200 | 0.0003260869 | 0 |
| DNA REPAIR | 1.4932768 | 142 | 0.0660994 | 0.005540166 |
| UNFOLDED PROTEIN RESPONSE | 1.3684303 | 108 | 0.11430663 | 0.040935673 |
| GLYCOLYSIS | 1.346378 | 141 | 0.11045363 | 0.025236594 |
| G2M CHECKPOINT | 1.2337347 | 195 | 0.21977922 | 0.046296295 |
| HYPOXIA | -1.3164258 | 130 | 0.11773413 | 0.04024768 |
| APOPTOSIS | -1.5006275 | 121 | 0.02761571 | 0.011128776 |
| IL2 STAT5 SIGNALING | -1.5871689 | 141 | 0.012231752 | 0 |
| APICAL SURFACE | -1.6423756 | 20 | 0.007078671 | 0.012567325 |
| IL6 JAK STAT3 SIGNALING | -1.6439723 | 53 | 0.007618738 | 0.0050083473 |
| INTERFERON ALPHA RESPONSE | -1.6528424 | 82 | 0.007266525 | 0 |
| APICAL JUNCTION | -1.7051502 | 104 | 0.004435164 | 0 |
| EPITHELIAL MESENCHYMAL TRANSITION | -1.7224233 | 67 | 0.004077788 | 0 |
| COAGULATION | -1.8201613 | 47 | 0.0017485606 | 0 |
| COMPLEMENT | -1.9678209 | 131 | 0 | 0 |
| ALLOGRAFT REJECTION | -2.0235777 | 135 | 0 | 0 |
| KRAS SIGNALING UP | -2.0399966 | 92 | 0 | 0 |
| INTERFERON GAMMA RESPONSE | -2.0819728 | 163 | 0 | 0 |
| TNFA SIGNALING VIA NFKB | -2.111887 | 146 | 0 | 0 |
| INFLAMMATORY RESPONSE | -2.260856 | 108 | 0 | 0 |

**Supplementary Table 11:** Enriched pathways in MEC-1 MYC-OE vs MYC-WT

#### Supplementary Figure legends

**Figure S1:** A. Kaplan–Meier plots showing the association between *MYC* expression and the time to first treatment (left) and overall survival (right) in 531 untreated CLL patients from the CLL-map portal. CLL patients were grouped according to MYC expression levels: $MYC^{low}$ (light purple line) and $MYC^{high}$ (dark purple line).

**Figure S2:** Generation of CRISPR/SAM-edited and shRNA/cDNA models. (A) Workflow of the generation of CRISPR/SAM-edited $MYC^{OE}$ PGA-1 and MEC-1 to introduce $MYC^{OE}$ (above) and the alternative models of PGA- 1 cell line to overexpress MYC by cDNA and inactivate *TP53* by shRNA (below). Western Blot (B), inmunofluoresence (C) and RTqPCR (D) showing $MYC^{OE}$ with 5 different sgRNAs in MEC-1 cell line. (E) Western Blot of *MYC* ant *TP53* of the shRNA/cDNA PGA-1 models.

**Figure S3:** $MYC^{OE}$ associated with higher proliferation rate in $TP53^{WT}$ cell lines. (A) Viability assay in shRNA/cDNA PGA-1 models. (B) Apoptosis assay in shRNA/cDNA PGA-1 models. % apoptosis (left graph) and representative annexin V/PI dot-plots (right panel) to determine apoptotic (annexin V + /PI +) cells. This experiment was made with two different replicates. (C) Cell cycle assay in shRNA/cDNA PGA-1 model. Representative cell cycle profiles using PI staining for each condition (up panel) and % values of different phases of cell cycle (down panel). This experiment was made with three different replicates. (D) *CCNE1* relative expression by RT-qPCR in CRISPR/SAM-edited PGA-1 model.

**Figure S4:** Different transcriptomic profiles when *MYC* is overexpressed in cellular models. PCA analysis of CRISPR/SAM-edited PGA-1 (A) and (B) in MEC-1. (C) *TP53* and *MYC* expression from RNA-seq data of CRISPR/SAM-edited cellular models. Legends: S1 represents sgRNA 19, S2 represents sgRNA 240 and S3 represents sgRNA 369

**Figure S5:** Other deregulated genes in $TP53^{WT}$ CLL patients and cellular models (A) Correlation analysis between *MYC* and *PDE4D (left)* and *ARHGAP32* (right) mRNA expression normalized in $TP53^{WT}$ CLL patients., Spearman's correlation was calculated in patients with detected expression in both genes using Log10 values for graphs visualization. (B-C) RT-qPCR in CRISPR/SAM-edited PGA-1 models showing the relative expression of deregulated genes that are commonly deregulated in common with $TP53^{WT}$ CLL patients (B) and other interesting genes associated with proliferation (C).

**Figure S6:** Drug sensitivity related with biological effects A) % cell viability of venetoclax (A-B), ibrutinib (C) drugs in CRISPR/SAM models. Cells were treated at different doses of venetoclax (1.56 and 0.781 μM), ibrutinib (1.561 μM) for 72 h. Cell viability was assessed with CCK-8, and surviving fraction is expressed relative to DMSO control. (D) % cell viability of different BTKi (acalabrutinib, zanabrutiniib and pirtobrutinib) at 20μM in *TP53*$^{KD}$-*MYC*$^{OE}$ PGA-1. (E) Dose–response curve of JQ1 treatment in shRNA/cDNA PGA-1 models. Cells were treated at different doses (0.098–3.125 μM) for 72 h. Cell viability was assessed with CCK-8, and surviving fraction is expressed relative to DMSO control. (F) Western blot MECOM in CRISPR/SAM-edited PGA-1 models. (G) MECOM relative expression by RT-qPCR in shRNA/cDNA PGA-1 models. (H) Enriched pathways related with unfolded protein response, G2/M checkpoint and DNA repair in CRISPR/SAM *MYC*$^{OE}$ MEC-1. (I) Dose–response curves of abemaciclib, talazoparib and belinostat in shRNA/cDNA PGA-1 models. Cells were treated at different doses (0.1–10 μM) for 72 h. Cell viability was assessed with CCK-8, and surviving fraction is expressed relative to DMSO control. (J) Dose-response curves of venetoclax (J) and ibrutinib (K) at increasing doses with STF-31 (19.5nM) for 72h in *TP53*$^{KD}$-*MYC*$^{OE}$ PGA-1 model.

**Supplementary Figure:**

**Figure S1:**

A

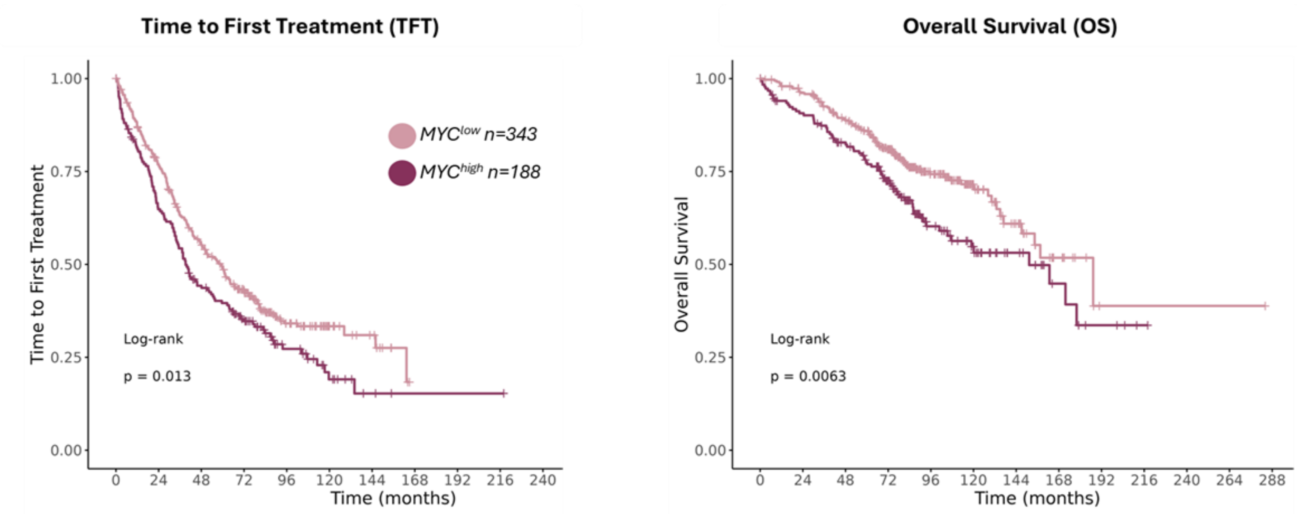

**Figure S2:**

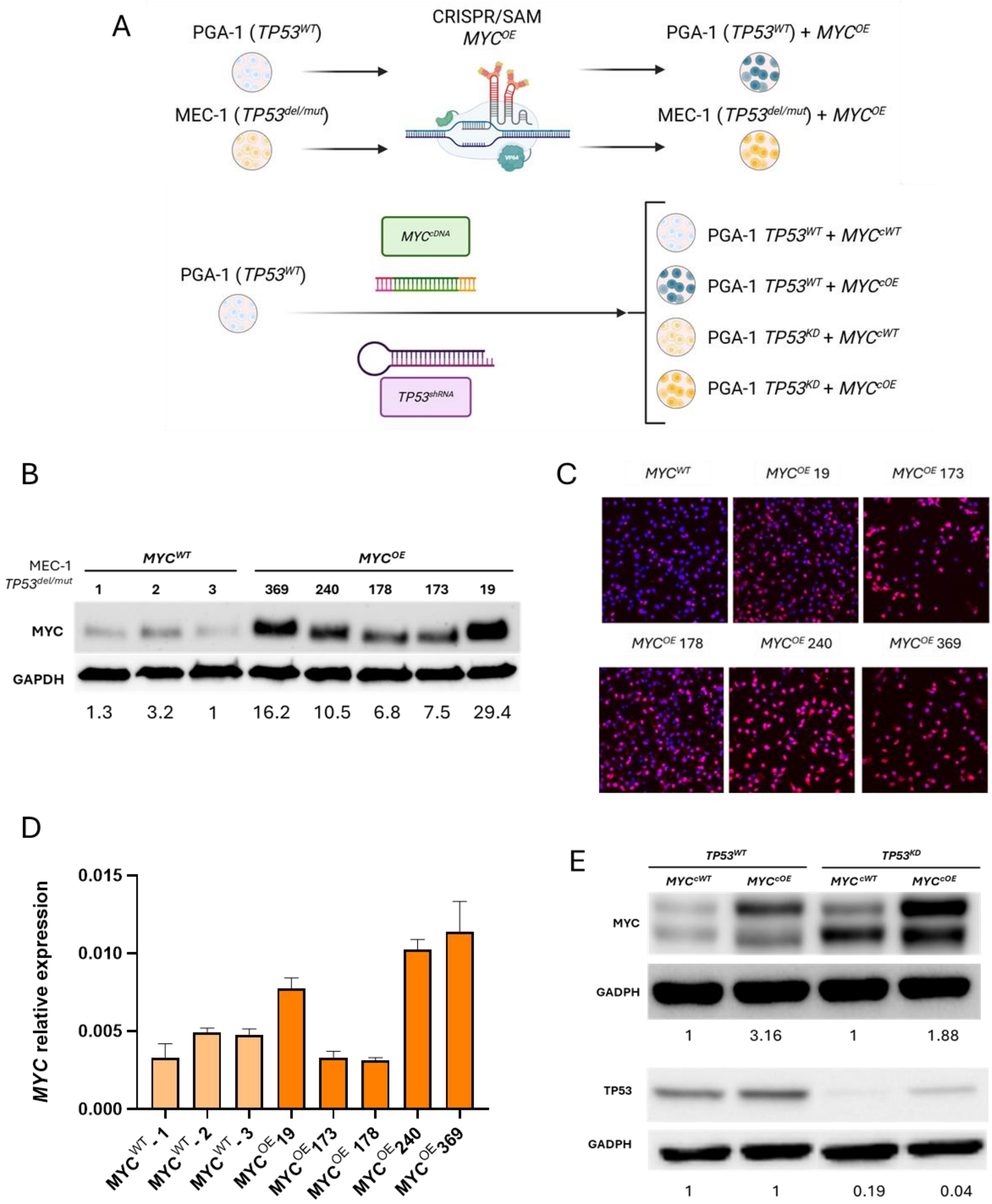
A
PGA-1 (TP53WT)
MEC-1 (TP53del/mut)
CRISPR/SAM
MYCOE
PGA-1 (TP53WT) + MYCOE
MEC-1 (TP53del/mut) + MYCOE
MYCcDNA
PGA-1 (TP53WT)
TP53shRNA
PGA-1 TP53WT + MYCcWT
PGA-1 TP53WT + MYCcOE
PGA-1 TP53KD + MYCcWT
PGA-1 TP53KD + MYCcOE
B
MEC-1
TP53del/mut
MYCWT
MYCOE
1 2 3 369 240 178 173 19
MYC
GAPDH
1.3 3.2 1 16.2 10.5 6.8 7.5 29.4
C
MYCWT
MYCOE 19
MYCOE 173
MYCOE 178
MYCOE 240
MYCOE 369
D
MYC relative expression
0.015
0.010
0.005
0.000
MYCWT_1
MYCWT_2
MYCWT_3
MYCOE19
MYCOE173
MYCOE 178
MYCOE240
MYCOE369
E
TP53WT
TP53KD
MYCcWT MYCcOE MYCcWT MYCcOE
MYC
GADPH
1 3.16 1 1.88
TP53
GADPH
1 1 0.19 0.04

**Figure S3:**

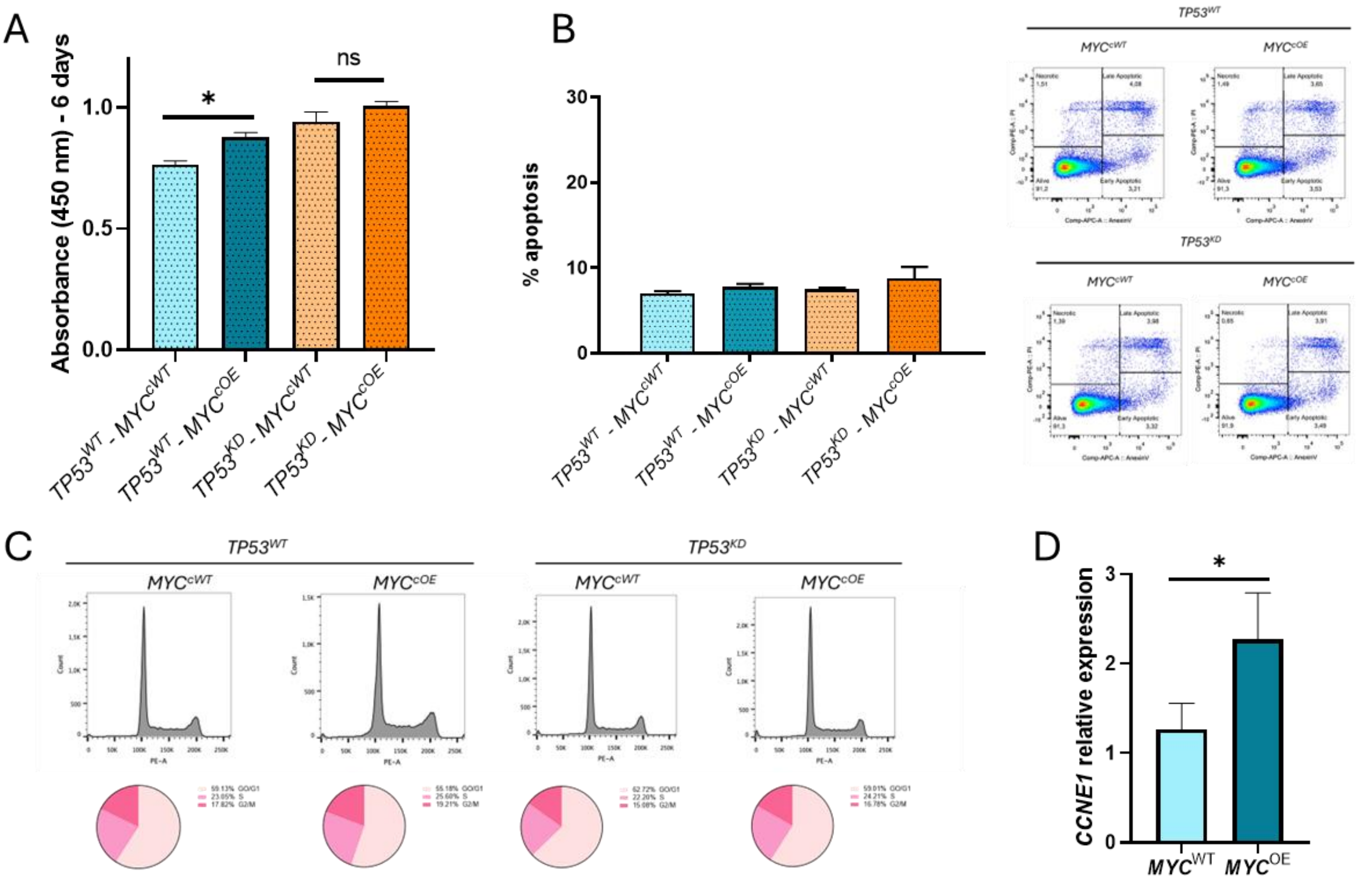
A
Absorbance (450 nm) - 6 days
*
ns
TP53^WT - MYC^cWT
TP53^WT - MYC^cOE
TP53^KD - MYC^cWT
TP53^KD - MYC^cOE
B
% apoptosis
TP53^WT
TP53^KD
MYC^cWT
MYC^cOE
C
TP53^WT
TP53^KD
MYC^cWT
MYC^cOE
D
CCNE1 relative expression
*
MYC^WT
MYC^OE

### Figure S4:

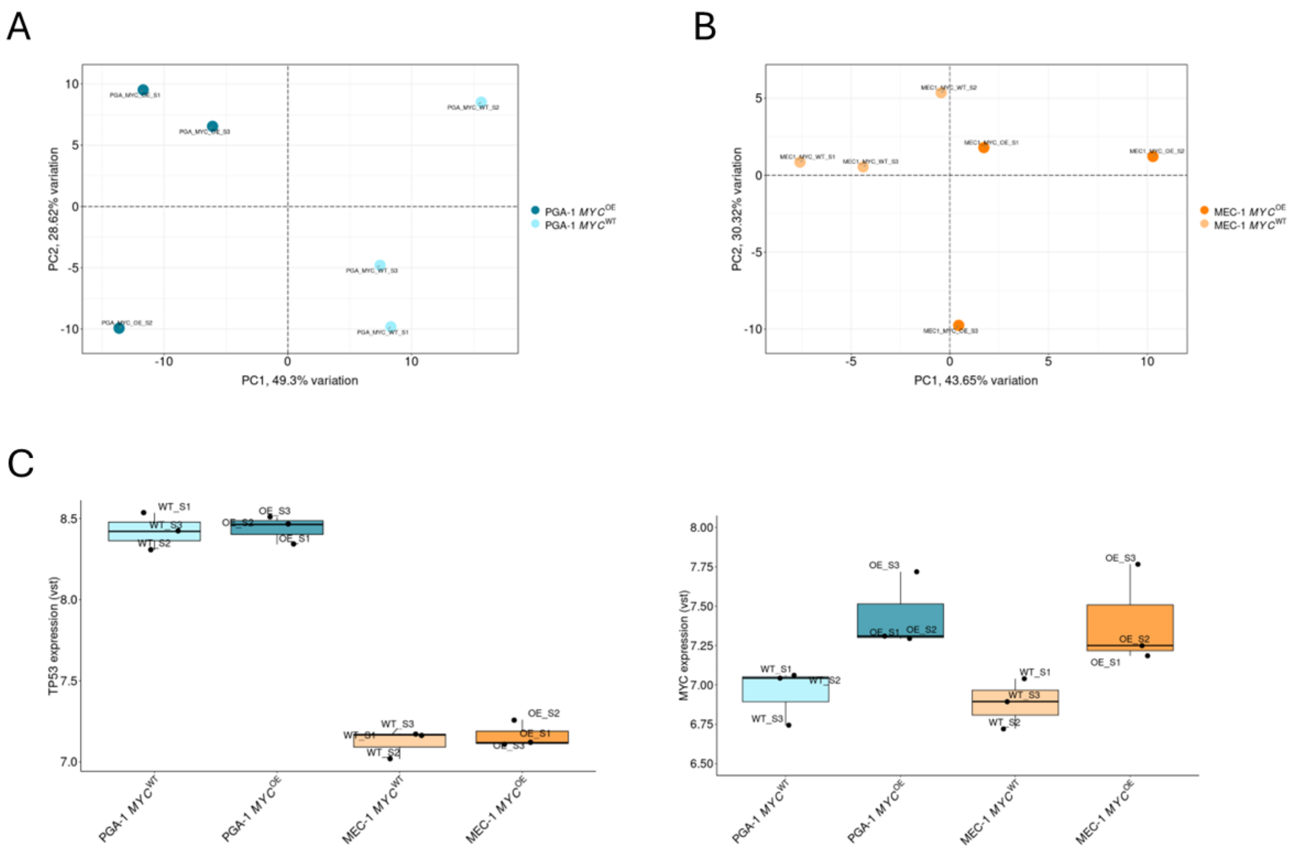

A
PC2, 28.62% variation
PC1, 49.3% variation
PGA-1 MYC^OE
PGA-1 MYC^WT
B
PC2, 30.32% variation
PC1, 43.65% variation
MEC-1 MYC^OE
MEC-1 MYC^WT
C
TP53 expression (vst)
MYC expression (vst)
PGA-1 MYC^WT
PGA-1 MYC^OE
MEC-1 MYC^WT
MEC-1 MYC^OE

### Figure S5:

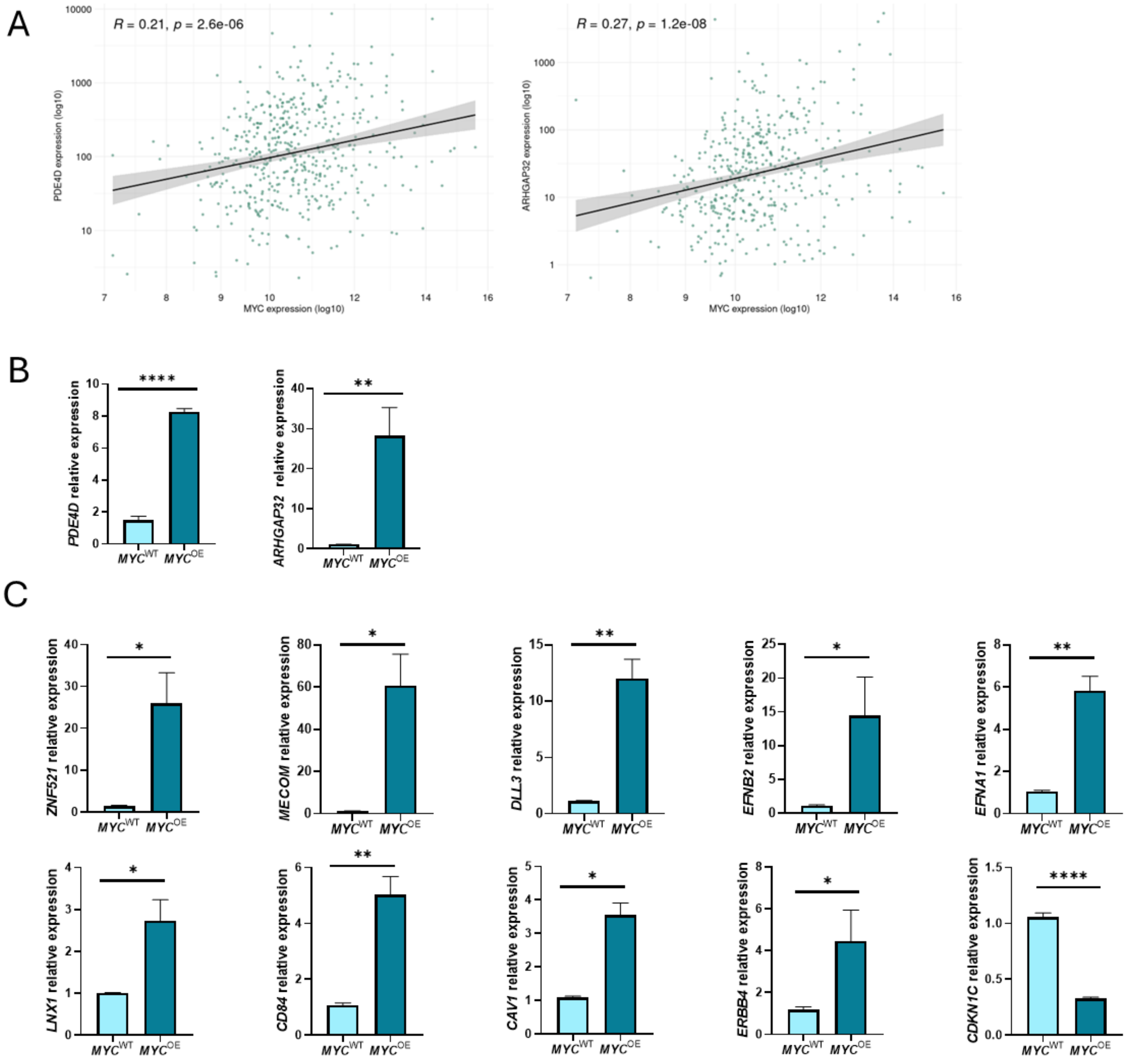

A
R = 0.21, p = 2.6e-06
PDE4D expression (log10)
MYC expression (log10)
R = 0.27, p = 1.2e-08
ARHGAP32 expression (log10)
MYC expression (log10)
B
PDE4D relative expression
ARHGAP32 relative expression
MYCWT MYCOE
C
ZNF521 relative expression
MECOM relative expression
DLL3 relative expression
EFNB2 relative expression
EFNA1 relative expression
LNX1 relative expression
CD84 relative expression
CAV1 relative expression
ERBB4 relative expression
CDKN1C relative expression

### Figure S6:

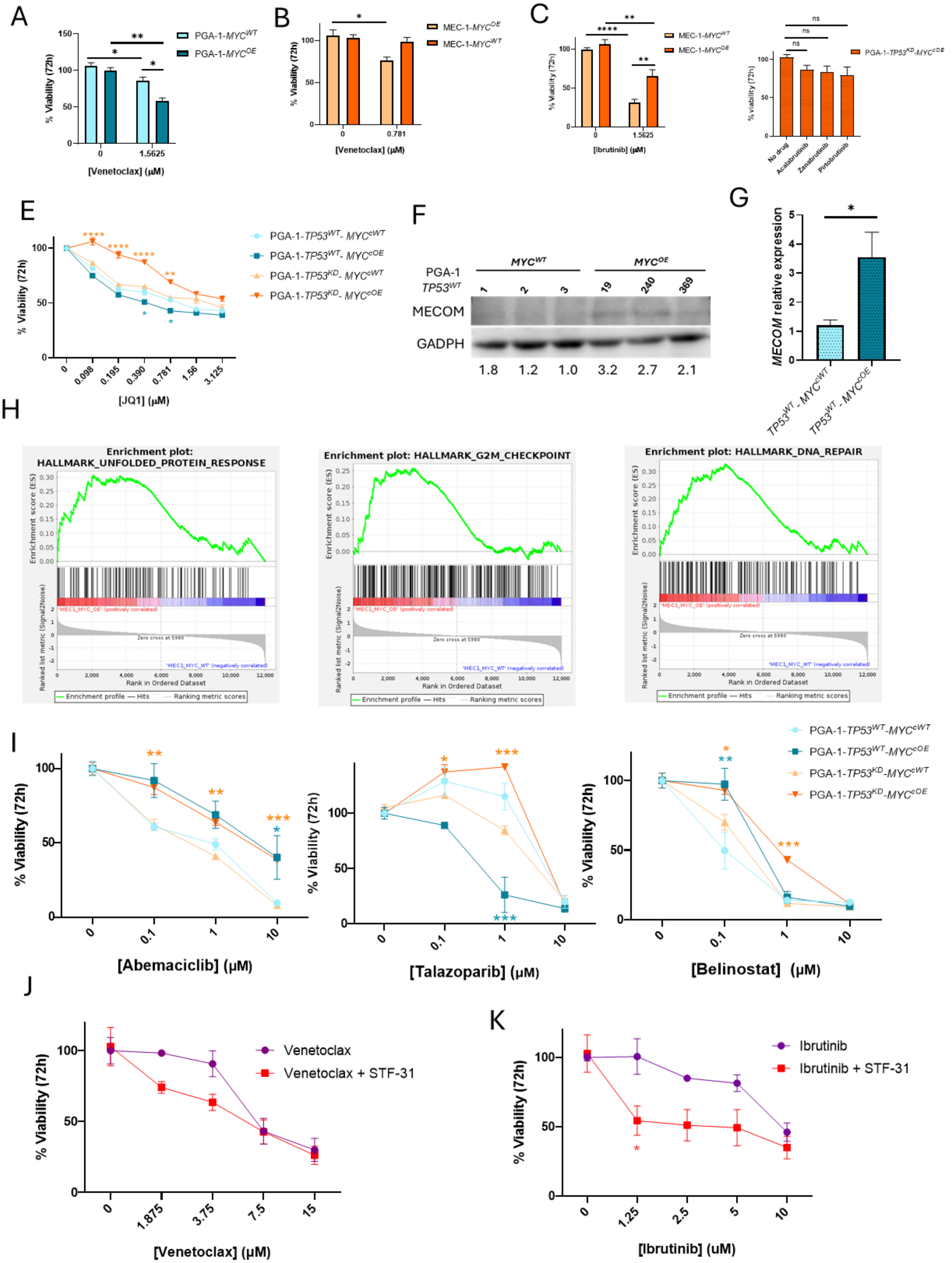

A
PGA-1-MYCWT
PGA-1-MYCOE
% Viability (72h)
[Venetoclax] (µM)
B
MEC-1-MYCOE
MEC-1-MYCWT
% Viability (72h)
[Venetoclax] (µM)
C
MEC-1-MYCWT
MEC-1-MYCOE
% Viability (72h)
[Ibrutinib] (µM)
PGA-1-TP53KD-MYCcOE
% viability (72h)
No drug
Acalabrutinib
Zanabrutinib
Pirtobrutinib
E
PGA-1-TP53WT- MYCcWT
PGA-1-TP53WT- MYCcOE
PGA-1-TP53KD- MYCcWT
PGA-1-TP53KD- MYCcOE
% Viability (72h)
[JQ1] (µM)
F
PGA-1 TP53WT
MYCWT
MYCOE
MECOM
GADPH
1.8 1.2 1.0 3.2 2.7 2.1
G
MECOM relative expression
TP53WT - MYCcWT
TP53WT - MYCcOE
H
Enrichment plot: HALLMARK_UNFOLDED_PROTEIN_RESPONSE
Enrichment plot: HALLMARK_G2M_CHECKPOINT
Enrichment plot: HALLMARK_DNA_REPAIR
Enrichment score (ES)
Ranked list metric (Signal2Noise)
Rank in Ordered Dataset
Zero cross at 5980
Enrichment profile — Hits — Ranking metric scores
I
PGA-1-TP53WT-MYCcWT
PGA-1-TP53WT-MYCcOE
PGA-1-TP53KD-MYCcWT
PGA-1-TP53KD-MYCcOE
% Viability (72h)
[Abemaciclib] (µM)
[Talazoparib] (µM)
[Belinostat] (µM)
J
Venetoclax
Venetoclax + STF-31
% Viability (72h)
[Venetoclax] (µM)
K
Ibrutinib
Ibrutinib + STF-31
% Viability (72h)
[Ibrutinib] (uM)
